# Stereo Slant Discrimination of Planar 3D Surfaces: Standard vs. Planar Cross-Correlation

**DOI:** 10.1101/2021.03.11.434881

**Authors:** Can Oluk, Kathryn Bonnen, Johannes Burge, Lawrence K. Cormack, Wilson S. Geisler

## Abstract

Binocular stereo cues are important for discriminating 3D surface orientation, especially at near distances. We devised a single-interval task where observers discriminated the slant of a densely textured planar test surface relative to a textured planar surround reference surface. Although surfaces were rendered with correct perspective, the stimuli were designed so that the binocular cues dominated performance. Slant discrimination performance was measured as a function of the reference slant and the level of uncorrelated white noise added to the test-plane images in the left and right eye. We compared human performance with an approximate ideal observer (planar cross correlation, PCC) and two sub-ideal observers. The PCC observer uses the image in one eye and back projection to predict the test image in the other eye for all possible slants, tilts, and distances. The estimated slant, tilt, and distance are determined by the prediction that most closely matches the measured image in the other eye. The first sub-ideal observer (local PCC, LPCC) applies planar cross correlation over local neighborhoods and then pools estimates across the test plane. The second sub-optimal observer (standard cross correlation, SCC), uses only positional disparity information. We find that the ideal observer (PCC) and the first sub-ideal observer (LPCC) outperform the second sub-ideal observer (SCC), demonstrating the benefits of structural disparities. We also find that all three model observers can account for human performance, if two free parameters are included: a fixed small level of internal estimation noise, and a fixed overall efficiency scalar on slant discriminability.

**Precis:** We measured human stereo slant discrimination thresholds for accurately-rendered textured surfaces designed so that performance is dominated by binocular-disparity cues. We compared human performance with an approximate ideal observer and two sub-ideal observers.

## Introduction

Estimating the 3D shape of our surroundings is essential for many everyday behaviors. The 3D shape at any point on a smooth surface can be closely approximated over a small neighborhood by a plane. Thus, the most local and fundamental measure of shape is local surface orientation. Local surface orientation is often specified in terms of slant and tilt (Stevens, 1983). Slant is the angle between the surface normal (the unit vector perpendicular to the surface) and the frontoparallel plane (Figure 1A). Tilt is the orientation of the vector formed by projection of the surface normal onto the frontoparallel plane (Figure 1B).

**Figure 1.**
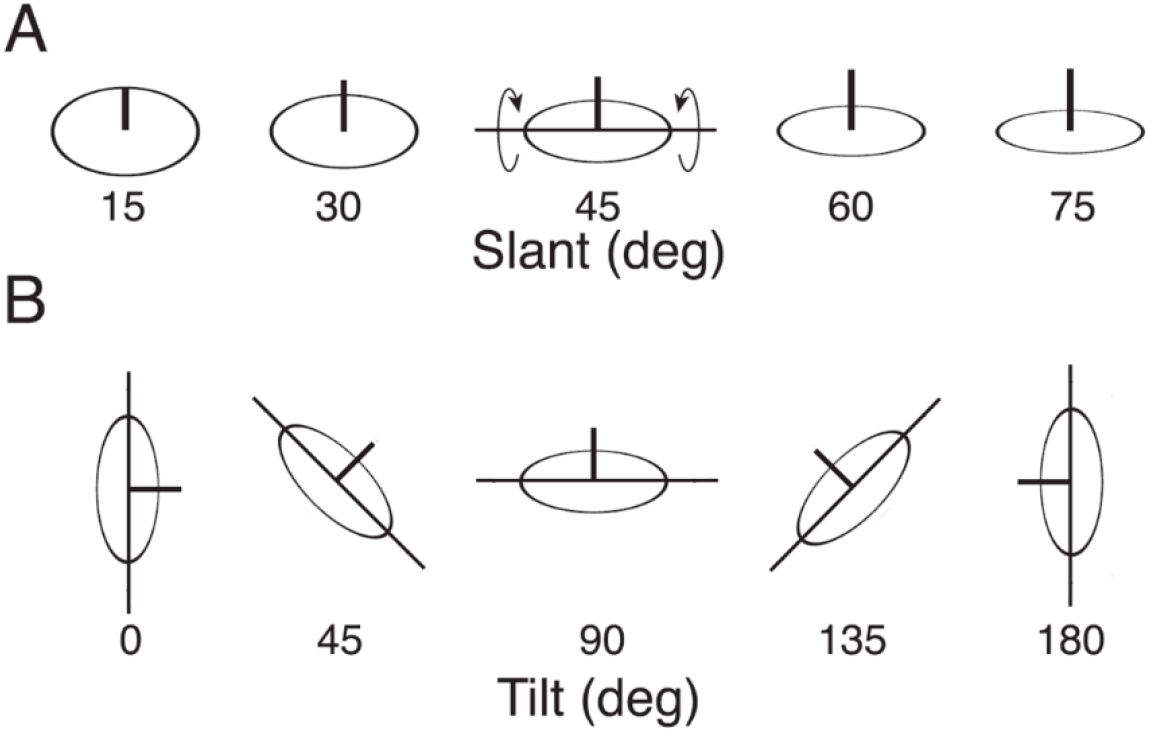
Definition of slant and tilt. **A** Slant is the angle between the surface normal (black vertical line segment) and the frontoparallel plane. Here the slant is varied while the tilt remains at 90°. **B** Tilt is the orientation of the vector formed by projection of the surface normal onto the frontoparallel plane. Here the tilt is varied while the slant remains at 45°.

A common view of 3D shape perception is that it begins with estimation of the local slants and tilts, which are then integrated into a representation of the 3D shape. Thus, not surprisingly, there have been a large number of studies directed at measuring and understanding the perception of 3D slant and tilt (e.g., see Howard & Rogers, 2012). Here we focus on perception of the 3D slant of planar surfaces.

Under natural conditions (without head or scene movement), the image information available for 3D slant estimation typically consists of the binocular cue of disparity (the differences between the images formed in the two eyes) together with various monocular cues (e.g., linear perspective). The primary goal of the current study was to measure slant discrimination under naturalistic conditions and to distinguish between specific hypotheses for how the human visual system estimates slant from binocular-disparity cues. A number of studies have measured slant-discrimination performance from binocular disparity using sparse random dot stereograms (Hibbard et al., 2002; Knill & Saunders, 2003; Hillis et al., 2004; Girshick & Banks, 2009; Burge et al., 2010). In most of these studies, the stimuli were presented in two temporal intervals.

In natural viewing, it is probably more typical for humans to be comparing the 3D orientations of surfaces that are densely textured, and that are located within the same scene at different distances (Burge et al., 2016; Kim & Burge, 2018; 2020). Here, we measured slant discrimination performance for surfaces that were textured with naturalistic noise (see Figure 3 in Methods). The two planar surfaces were presented in a single-interval task, where the smaller test surface was in front of a surrounding reference surface by a distance that varied randomly from trial-trial-trial by a small amount. The stimuli were accurately rendered, and hence contained both monocular and binocular cues to surface orientation. To reduce the usefulness of the monocular cues, the texture contained few regularities and the shape (i.e., silhouette) of the test surface was jittered (see Methods). A control experiment confirmed that the performance of our subjects was completely dominated by the binocular cues (see Results). This allowed us to focus on models of slant discrimination from binocular disparity. Specifically, we derived the approximate ideal observer for slant discrimination of planar surfaces and compared its performance, and those of various sub-ideal observers, with human performance. To facilitate comparison of human and model-observer performance, the slant discrimination thresholds were measured with various levels of uncorrelated white noise samples added to the left- and right-eye test regions (see Figure 4).

Ideal-observer models reveal the fundamental computational principles of the task, set a proper benchmark against which to compare human performance, and can be used to evaluate the effectiveness of heuristic (sub-optimal) mechanisms (Green & Swets, 1966; Geisler, 2011; Burge, 2020). To specify the approximate ideal observer here, we assumed that the geometrical relationship between the two eyes is known, and that the two images are projections of a single patch of planar surface at an unknown distance and with an unknown 3D orientation. The choice of coordinate system does not impact the discrimination performance of the ideal or sub-ideal observers considered here (see Appendix). For simplicity we assume a head-centered coordinate system where the optic axes of the left and right eyes are parallel to the head-centered depth axis, and where the left and right images are projections onto a cyclopean image plane located a fixed distance from the nodal points of the two eyes (see Figure 2A & 2B). In the Appendix we consider cases in which the eyes are not in primary position (i.e., not parallel to the cyclopean axis), and in which the projection is onto a spherical surface in each eye. These cases are important when considering specific hypotheses for the underlying anatomy and neurophysiology.

**Figure 2.**
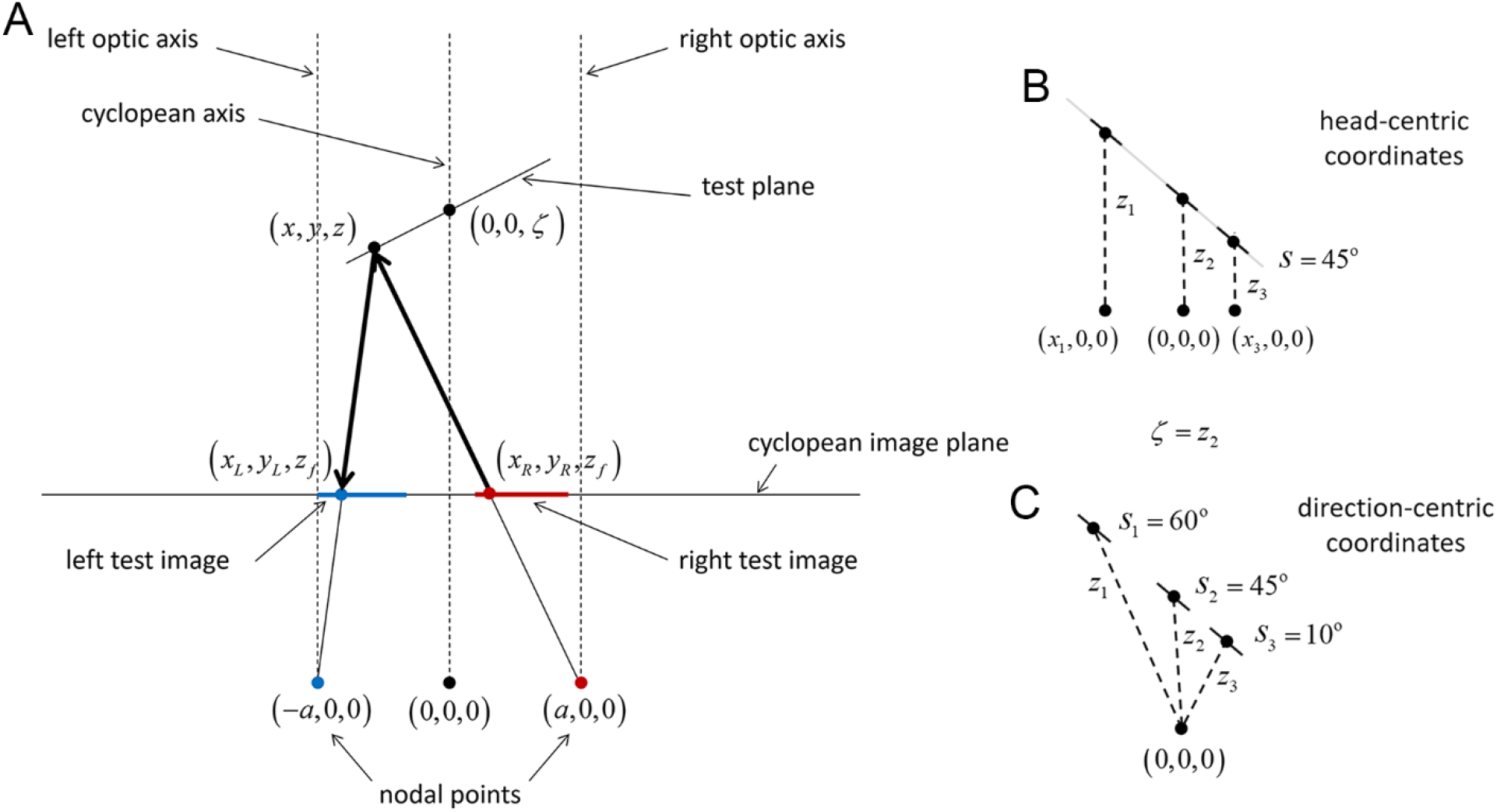
Schematic of the viewing and imaging geometry used for determining ideal and sub-ideal observer performance. **A**. The test plane is a planar surface whose distance *ζ* and slant *s* are defined with respect to the cyclopean axis. The left (blue) and right (red) images of the test plane are formed in the cyclopean image plane by perspective projection for nodal points separated by an interocular distance of 2*a*. The ideal observer uses planar cross correlation (PCC). For each possible slant and distance of the test plane, the observer predicts the left test image from the right test image (or vice versa) by back projection and then forward projection; for example, the predicted gray level at (*x*_*L*_, *y*_*L*_) is the gray level at (*x*_*R*_, *y*_*R*_). The estimated slant 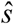 and distance 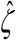 of the test plane are the values that give the smallest prediction error. **B**. In head-centric coordinates all points in the test plane have the same slant. The distance *ζ* of the test plane is the intercept of the plane, containing the test plane, with the cyclopean axis. The distance *z*_*i*_ of individual points in the test plane varies with location in the cyclopean image plane. **C**. In direction-centric coordinates the slant and distance are in general different for different points in the test plane. Model performance is the same for the two coordinate systems, but for simplicity we use head-centric coordinates.

For every possible slant and distance of the surface, the ideal observer computes the predicted image in one eye given the image observed in the other eye and the rules of forward- and back-projection. The estimated slant is the slant value from the slant and distance pair that gives the smallest prediction error. We will call this optimal model of slant and distance estimation the “planar cross correlation” (PCC) model (see Methods and Appendix). This observer is optimal because it uses all of the available geometric information given planar surfaces. By generating predictions via back projection for every possible distance and slant, the PCC observer is considering exactly the set of possible differences that can exist between the left and right images for a given planar surface. It then picks the distance and slant that best explains the difference between the two images.

Figure 3 illustrates this mapping for the more general case of an arbitrary 3D surface orientation (see also Appendix Figure A1). Figure 3B shows an image region in the right eye (red square) and the back projection to image plane for the left eye (blue trapezoid), for the correct distance and surface orientation (60° slant and 45° tilt). Figure 3C demonstrates this mapping with the stereo-image pair of a blurred checkerboard surface.

**Figure 3.**
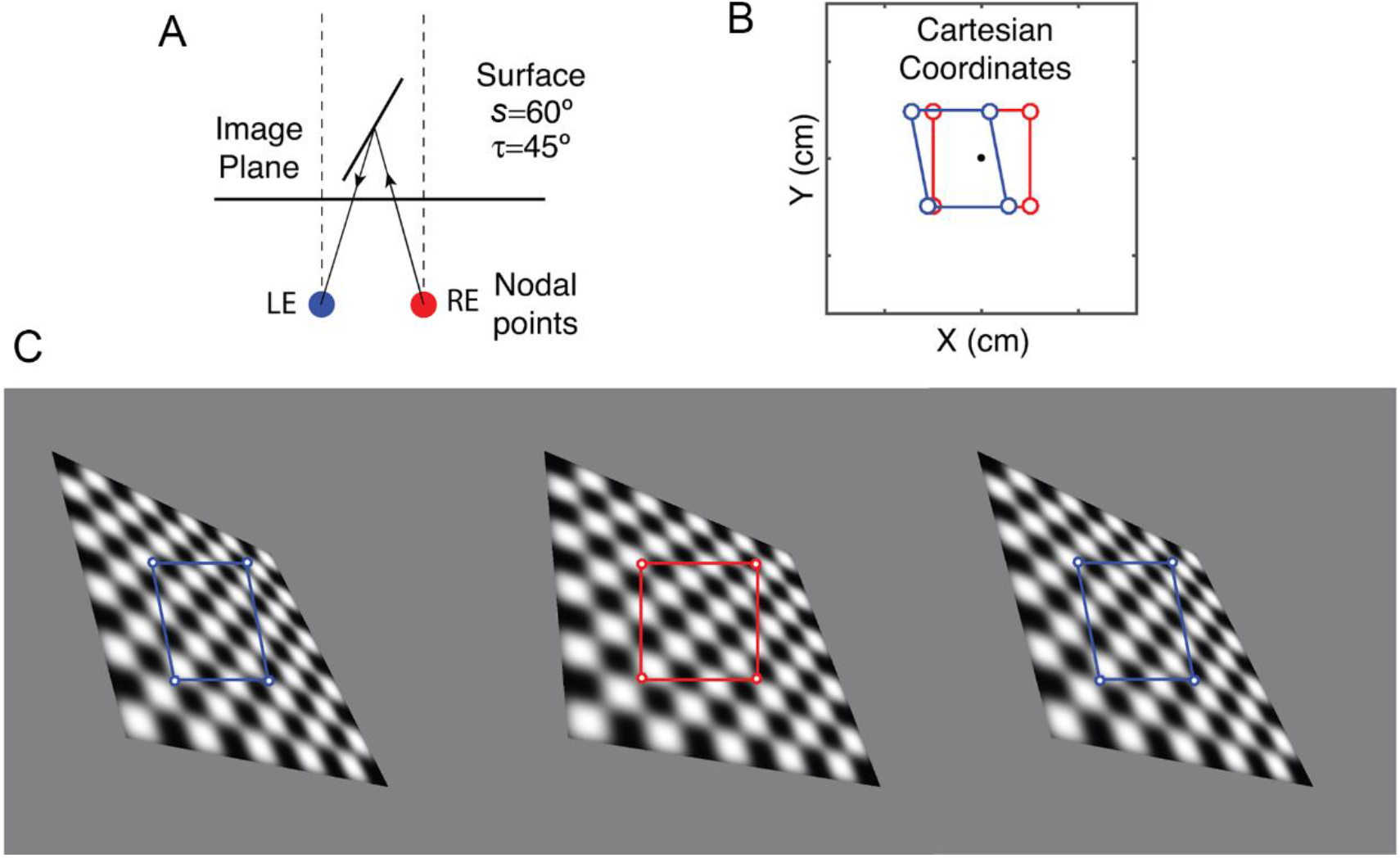
Image differences in Cartesian coordinates. **A**. Viewing geometry. **B**. A square image patch in the right eye view (red) and the back projection of its boundary to the left eye (blue), in Cartesian coordinates centered on the cyclopean eye. C. Stereo pair of surfaces and patches in each eye. Fuse images to see a surface with a slant of 60° and a tilt of 45° (crossed pair: left and center, uncrossed pair: center and right). Notice that when fused the red and blue boundaries coincide.

In general, human performance deviates from optimal performance. A principled approach for generating plausible sub-optimal models is to replace one or more of the optimal computations with simpler more biologically-plausible computations, to incorporate internal noise, and/or to incorporate other plausible biological limitations (e.g., response nonlinearities, foveation, etc.).

One simplifying, and more biologically plausible, computation is to perform planar cross correlation locally, and then combine the local slant and distance estimates over the whole test region (Jones & Malik, 1992; Super & Klarquist, 1997; see also Wildes, 1991). We will call this the “local planar cross correlation” (LPCC) model.

The most simplifying assumption is that the structured patterns of disparity produced by slanted surfaces (e.g., disparity patterns that are structured in orientation and scale) are not used directly to estimate 3D surface orientation. This assumption is made by most models of human stereo vision. Here we implement this simpler hypothesis by computing disparity at each location using “standard cross correlation” (SCC) (Tyler & Julesz, 1978; Cormack et al, 1991; Banks et al., 2004). The estimated slant is then computed by combining the distances specified by the disparities. Formally, this SCC model is a special case of LPCC model where the local slant is assumed to be zero (see Methods and Appendix). The SCC model uses only local disparities in position. Models based on standard cross-correlation have been successful in accounting for many aspects of human disparity discrimination (Banks et al., 2004; Cormack et al., 1991; Filippini & Banks, 2009; Tyler & Julesz, 1978), and in explaining the response properties of disparity-selective neurons in visual cortex (Ohzawa et al., 1990; DeAngelis et al., 1991; Ohzawa et al., 1997; Cumming & DeAngelis, 2001).

It has long been known that introducing an orientation or scale difference between the left and right images can produce vivid perceptions of surface slant (Wheatstone, 1838; Ogle, 1938; Blakemore, 1970a). These results seem to suggest that structured patterns of disparity in orientation and scale are directly exploited by the visual system to estimate 3D surface orientation. However, these structured disparity patterns are confounded with local horizontal disparities. Thus, it has been difficult to rule out the hypothesis that horizontal disparities are computed first and then later combined to determine 3D surface orientation (Fiorentini & Maffei, 1971; Wilson, 1976; Mitchison & McKee, 1990; Cagenello & Rogers, 1993; Halpern et al., 1996; Greenwald & Knill, 2009).

In the current study, we measured slant discrimination thresholds for the human and model observers as a function of reference slant and the level of white noise, uncorrelated samples of which were added to each eye’s image. As expected, we found that the PCC model had the lowest thresholds, followed by the LPCC model, the SCC model, and finally the human subjects. All three models capture the qualitative trends in the human thresholds, but none provide good quantitative predictions of the trends, even when their average sensitivities (*d′* values) are scaled by an arbitrary efficiency parameter. However, if we include another known factor, internal noise, all three models make good quantitative predictions. Specifically, we include a fixed low level of internal estimation noise. With this internal noise, the predictions are equally good for all three human observers (although each observer requires a different overall efficiency scalar). We also measured depth discrimination, in addition to slant discrimination, with the same stimuli and found that there was a trend for human observers to be more efficient (relative to ideal) at slant discrimination than at depth discrimination. We find that the PCC and LPCC computations are more accurate at slant estimation than the SCC computation, and thus there should have been evolutionary pressure to incorporate equivalent computations into the early visual system.

## Methods

### Subjects

Three experienced psychophysical observers (two males and one female) served as subjects. They each had normal or corrected to normal spatial and stereo acuity. Written informed consent was obtained for all observers in accordance with The University of Texas at Austin Institutional Review Board.

### Apparatus

Stimuli were presented using a Planar PX2611W stereoscopic display (Planar Systems, Beaverton, OR). This display consists of two monitors with orthogonal linear polarization relative to each other, separated by a polarization-preserving beam splitter. Subjects wore passive linearizing filters to view binocular stereo stimuli. In all experiments, subjects used a forehead rest to maintain constant viewing distance. Each monitor was gamma-corrected to produce a linear relationship between pixel values and output luminance. Luminance was measured with a photometer (PR 655, Photo Research; Syracuse, NY) through the beam splitter and a polarizing lens. The background luminance of the two monitors was 46.73 and 52.04 cd/m^2^. All experiments and analyses were done using custom code written in MATLAB using the Psychophysics Toolbox (Brainard, 1997; Pelli, 1997).

### Stimuli

The stimuli (Figure 4) were stereo images centered on a display screen located at a distance of 100 cm from the eyes. The stimulus consisted of two surfaces: a relatively large rectangular surround reference plane (with an angular central height of 9°), and a smaller central test plane (see Figure 4A). The slants of both the reference and test planes were varied; the tilts of the reference and test planes were always zero (i.e., both surfaces were slanted about a vertical axis). In 3D space, the rectangular reference plane had a fixed width (23 cm) and height (16 cm), and the center of the reference plane was located at a distance of 102 cm (i.e., 2 cm behind the display screen). There was a “window” (hole) in the reference plane that subtended 4.2° × 3.1° in the right eye, independent of the slant of the reference plane. Thus, the window size varied in the left eye when the slant of the reference plane was varied. The test plane was centered in the window of the reference plane. On each trial, an independent sample of spatial Gaussian white noise was added to the window regions of the left- and right-eye images. The reference slant and additive noise contrast were parametrically varied: reference slants = 0°, ± 12.5°, ±°; 25°, ±50°; root-mean-square (RMS) noise contrasts = 5%, 17.5%, 34%. The slant of the test plane was varied to obtain psychometric functions for slant discrimination (see below). The stereo images were rendered assuming an interocular distance of 6.5 cm (the interocular distances of the three subjects are 6.5, 6.5, and 6.2 cm).

**Figure 4.**
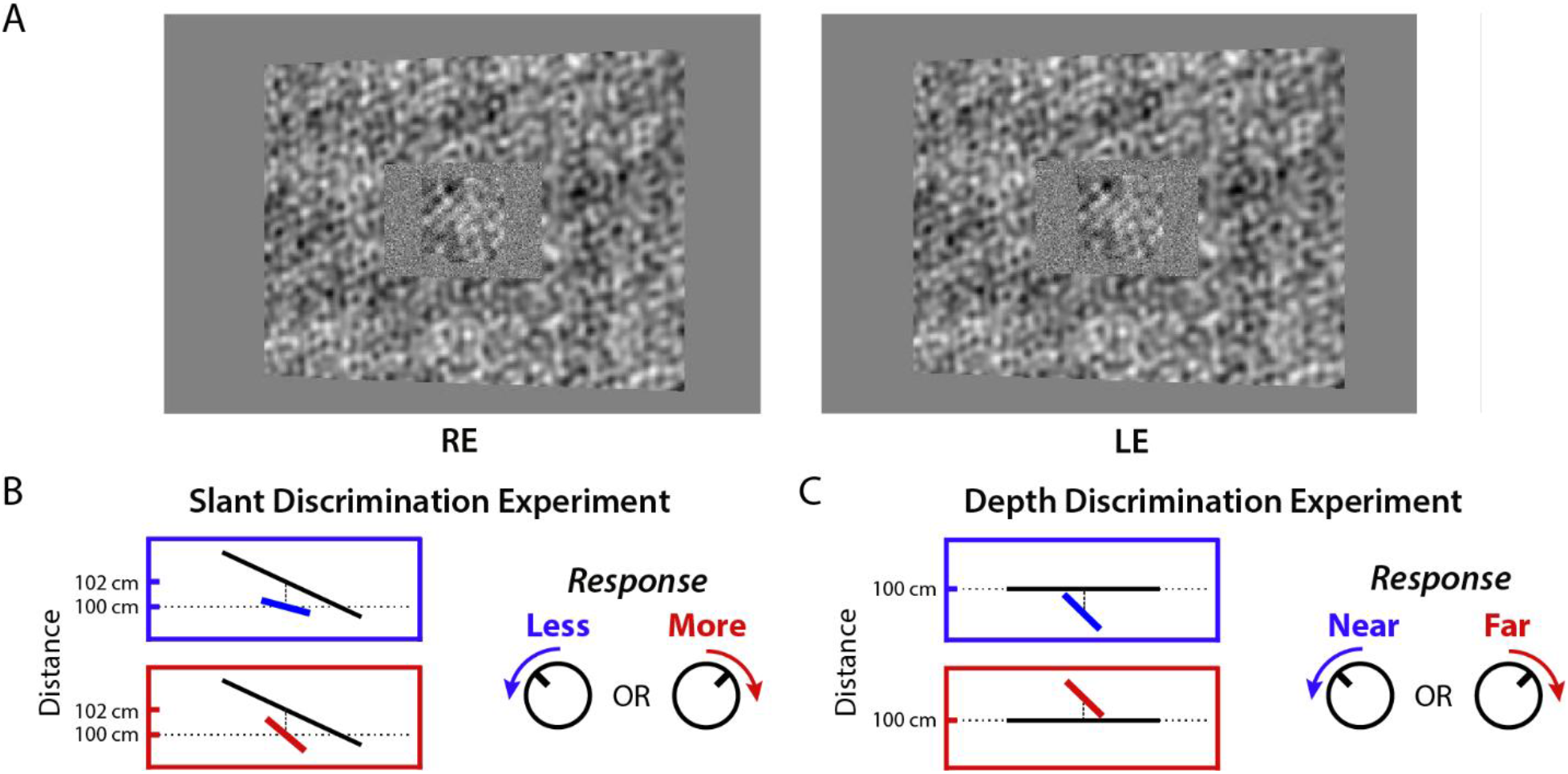
Stimuli and task in the slant and depth discrimination experiments. **A**. Example binocular stimulus (crossed). The actual stimuli were presented in a stereo rig where orthogonally-polarized left- and right-eye images alternated at 60 Hz, and were viewed through polarization-selective filters. The rectangular reference plane was densely textured, had a central window/hole, and was rendered at a distance of 102 cm. The central test plane was a trapezoid jittered in distance and aspect ratio to reduce the usability of monocular perspective cues. On each trial an independent sample of white noise was added to the window region in the left- and right-eye images. **B**. As illustrated in a top-down view, the subjects’ task was to judge whether the central test plane was more or less slanted that the reference plane. Subjects had unlimited viewing time and responded by rotating a knob clockwise or counter clockwise. **C**. Top down view of the depth discrimination experiment. The reference plane was frontoparallel and the subject judged whether the slanted test plane was near or far.

When a planar surface is slanted, monocular slant cues are created due to perspective projection. Our main interest here was in the stereo cues, and thus steps were taken to reduce the usefulness of the monocular cues. The effectiveness of these steps was confirmed with a monocular control experiment (see below). First, we randomly varied the distance of the test plane from 99-101 cm across trials, thereby jittering its retinal-image size. Second, we fixed the width and central height of the test plane in the right eye to 2.2 ° of visual angle and jittered the ratio of the left and right edge heights of the test plane. This jitter creates 3D test plane surfaces that are trapezoidal in shape. The average edge-height ratio was set equal to what would be expected for a rectangle having the same slant as the reference plane. The range of the random jitter around this ratio matched the range of ratios expected for a rectangular test plane varying over the range of test slants used to measure the psychometric function with the steepest slant (50°). This procedure strongly reduces the usefulness of monocular shape cues to slant estimation, while introducing a relatively small amount of jitter. Even with the jitter, some monocular cues for slant remained because the surface texture in the test plane was accurately rendered.

The texture of the planes was generated by summing 59 sinewaves having spatial frequencies of constant amplitude ranging between 0.1 and 3 cycles per degree (cpd) in steps 0.05 cpd, with the phase and orientation of each sinewave randomly sampled from a uniform distribution over all possible values (0 to π). The advantage of creating textures by summing sinewaves is that it is possible to avoid interpolation artifacts. For each pixel location, for each eye, there is an exact formula for the real-valued gray level for each sine-wave component on any 3D planar surface. Given these values, we then then summed the gray levels for all the components to obtain the exact real-valued gray level at each image pixel location, and then be gamma-compressed and appropriately quantized them for presentation. The textures on the reference and test planes were generated separately, and both had an average RMS contrast of 14.7% before addition of the uncorrelated white noise.

One difference between the task used here and tasks used in previous investigations of slant discrimination is that the task is arguably more typical of natural conditions. Typically, the two surfaces whose slants are being compared are presented in two different temporal intervals (Knill & Saunders, 2003; Hillis et al., 2004; Girshick & Banks, 2009). Under natural conditions, surfaces that are being compared are often at different distances and are often viewed simultaneously. Also, in order to isolate binocular disparity cues, many previous studies have used sparse random-dot stereograms. However, most natural surfaces have a dense irregular texture (e.g., tree bark).

### Procedure

The experiment was performed under free-viewing conditions without a fixation point, although a chin and head rest were used to fix the head position. Observers were asked to indicate, in a forced choice task, whether the test plane was more or less slanted than the reference plane. To indicate their decision, observers simply turned a knob (PowerMate wireless controller Griffin Technology, Irvine, CA) in the direction in which the test plane was rotated relative to the reference plane. Observers found this method of response much more intuitive than a keypress. The stimulus was present until the observer made a response. The average trial duration was 4.7 seconds. The next stimulus appeared after a 1.5 s blank interval.

Each observer completed eight experimental sessions, where the magnitude of the reference slant was held fixed at one of the four values (± 0°, ± 12.5°, ± 25° and ± 50°). Each session consisted of four blocks of trials. The first block was a practice block where the number of trials was half that of the other blocks and with feedback given on each trial. Practice blocks were not included in the data analysis. No feedback was given in the remaining three experimental blocks (160 trials per block). In each of these three blocks the noise contrast was fixed at one the three values (5%, 17.5%, 34%). Within a session, the order of these blocks was either ascending or descending, and in the later repeat of that session the order of the noise contrasts was reversed. All 160 trials in a block had a fixed reference slant magnitude, but the sign of the reference slant was different for the first and second halves of the trials in the block (e.g., 25° in the first half and −25° in the second half). We combined all trials having the same magnitude of slant and hence there was a total of 360 trials per condition.

Within each block, a psychometric function was measured by varying the slant of the test plane relative to the fixed reference plane. There were eight levels of slant per psychometric function presented in a random order. The texture of the test plane was different on each trial. The texture of the reference plane was different in each block. All observers made judgments for the same stimuli, with a different random order for each observer.

### Control Experiments

We ran two control experiments in addition to the main experiment, . The first control experiment was a slant-discrimination experiment where the viewing was monocular. Measurements (a total of 160 trials per white-noise level) were only made for the steepest slant, because those stimuli contain the steepest texture gradient, and thus the most reliable monocular information (Knill, 1998; Hillis et al., 2004). The stimuli were constructed using the same rules as those in the main experiment, but observers viewed the stimuli with a patch over the right eye. For all noise levels and slant differences between test and reference plane used in the actual experiment, the three observers performed at chance.

In the second control experiment, the three observers were asked to discriminate the depth rather than the slant of the test plane relative to the reference plane (see Figure 3C). The reference plane was given a 0° slant and was rendered at a distance of 100 cm. The slant of the test plane was fixed at either −44° or 44° so that the near and far edges of the test plane in 3D space were approximately 2 cm in front and behind the reference plane. The test plane was slanted so that the disparity pedestal values (difference in baseline disparity from zero disparity) were similar to those in the main experiment. In depth discrimination experiments it is known that threshold tends to increase with the magnitude of the disparity pedestal (Blakemore, 1970b; Schumer & Julesz, 1984; Badcock & Schor, 1985; Stevenson et al., 1992). Thus, for comparing human efficiency in the two tasks it is best to keep the average pedestal disparities similar.

Depth discrimination of the test plane was measured as function of noise contrast. Each participant completed two sessions, one in increasing order of noise contrast and one in decreasing order. Each session consisted of three pairs of blocks, one pair for each noise contrast. The first block in each pair had half the number of trials, and feedback was provided. There was no feedback in the second block of each pair. There were 80 trials in each no-feedback block. The data from the feedback blocks (40 trials) was not used in the analysis. To measure psychometric functions, the depth difference between the reference and test planes varied within the block. There were eight depth difference levels presented in random order. The observers’ task was to report whether the center of the test plane was closer or further than the reference plane. As in the main experiment, different random textures were generated for each trial, and all observers made judgments for the same stimuli. The only difference was that the order of the stimuli was randomly different for each observer.

### Analysis Methods

For non-zero reference slants the response data was the percentage of “more-slanted” responses as function of the angular difference between test and reference planes. For the zero reference slant (frontoparallel reference plane) the response data was the percentage of “slanted-left” responses as a function of test slant. For each combination of reference-slant magnitude and noise contrast, we first merged the psychometric data for the two reference-slant orientations having the same magnitude (e.g., 50° and −50°). The psychometric data for each condition and subject was then fitted with a cumulative Gaussian function using maximum likelihood,

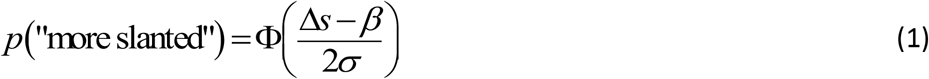

where Δ*s* is the test slant minus the reference slant, *σ* is the standard deviation parameter, and *β* is the bias parameter. The bias parameter corresponds to the 50% point of the psychometric function, and the value of the standard deviation was defined to be the threshold. Note that the discriminability, *d′* , equals Δ*s*/*σ*. We define threshold to be the value of Δ*s* for which *d′* =1.

Not surprisingly, given the well-known individual differences in stereo acuity (Coutant & Westheimer, 1993; Bohr & Read, 2013), there were substantial differences in overall performance level between the three observers. However, the three pair-wise correlations between participants’ thresholds were quite high (0.967, 0.973, 0.962). Therefore, to better see the average trends, we scaled the thresholds of two of the observers so that their average threshold was the same as the average threshold of the intermediate performing observer. The scale factor for the more sensitive observer was 0.68 and for the less sensitive observer was 2.01. These same scale factors were also used to scale the biases.

The control experiments were analyzed using the same procedures. However, for the control experiment on depth discrimination, the differences in overall performance between observers were not as large, so the data was averaged without scaling.

## Models

We consider an approximate ideal observer and several suboptimal observers. In all cases, the input to the model is the pair of left and right images produced by the test plane. For mathematical convenience we assume a zero vergence angle and a cyclopean image plane that is perpendicular to the optic axes (Figure 2). More general cases are described in the Appendix. However, we note that as long as the orientations of the eyes and the shape of the imaging surfaces are known, then the information available for estimating surface slant remains largely invariant. Thus, the performance of the models for the geometry in Figure 2 is quite general. Of course, the specific computations of the models depend on the orientations of the eyes and shape of the imaging surfaces.

### Approximate Ideal Observer: Planar Cross Correlation (PCC)

Derivation of the exact ideal observer would start with a statistical description of the 3D surfaces, a deterministic description of the imaging geometry, a statistical description of the uncorrelated noise added to the two images, a description of the prior probability on surface slant and distance, and a description of the cost function for the task. The derivation would then involve solving for the specific computations that minimize the cost function. The exact derivation was not practical for our particular stimuli, but given our past experience with similar models, we think that the computations described below should be a close approximation to ideal.

As mentioned earlier, we assume that the geometrical relationship between the two eyes is completely known and fixed, and that the two test images are projections of a single planar surface at an unknown intercept distance and slant. In this case, it is optimal to compute the predicted image in one of the eyes given the image in the other eye. Specifically, for every possible distance and slant of the surface, the predicted left-eye image is computed from the right-eye image by projecting points in the right-eye image to the corresponding point in left-eye image (see Figures 2 and 3). The predicted real-valued location in the left image 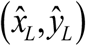, given a location (*x*_*R*_, *y*_*R*_) in the right image, is given by the equations in Appendix Figure A1. The predicted gray level at the predicted location in the left image, is equal to the gray level in the right image at the original location: 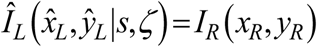. The predicted left-eye image is then compared to the actual left-eye image. The estimated slant is the slant value, from the slant and z-intercept pair, that gives the smallest prediction error (mean squared error):

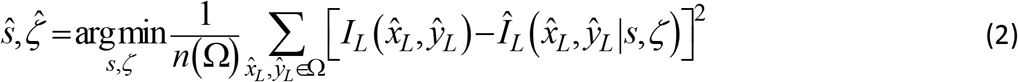

where Ω is the set of predicted left-eye image coordinates, *n*(Ω) size of the set, and 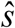, 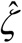 are the estimated slant and intercept distance of the plane.

Predictions were also generated for a few conditions using a typical normalized correlation error measure. It performs very similarly, but is computationally much slower. Also, for a simple absolute disparity estimation task (not slant estimation), we found that these two measures perform similarly and well-approximate the exact ideal observer for that task (Oluk & Geisler,2020).

The display was designed so that the image of the test plane in the right-eye image always had the same width (2.23°), which is assumed to be known by the model observers. To further simplify computations, the predictions were generated for a fixed height (2.12°) in the right-eye image, which corresponds to the minimum height produced in the experiments. This leaves out ~5% of the informative pixels, but has a minor effect on the predictions.

The test- and reference-plane textures were a sum of random sine waves having a maximum spatial frequency of 3 cpd, before rotation about the vertical axis. The image spatial frequencies in the test and reference plane can be higher because of the surface slant. In the experiment, uncorrelated white noise was added to the left- and right-eye test-plane regions. The ideal observer must pre-filter the stimuli before comparing the left- and right-eye images; that is, it must filter out spatial-frequencies in the added white noise that do not overlap with frequencies present in the signal. To approximate the optimal prefiltering, we carried out a preliminary analysis where we measured PCC performance for various values of the cutoff frequency of a low-pass filter that ramped to zero over a span of 1 cpd. This was done separately for each level of uncorrelated noise. We found that the optimal cutoff frequencies for the 5%, 17.5% and 34% noise contrasts are 16, 8, and 5 cpd, respectively. Variation in these cutoff frequencies by 1-2 cpd had little effect on performance. We used this same prefiltering for all model observers.

### Local Planar Cross Correlation (LPCC)

It is not biologically plausible that planar cross correction is computed in one step over the entire test region (especially if the test region were larger than the current 2.13° width). Thus, we also consider a sub-optimal version where planar cross correlation is computed over square right-eye patches of some given width *w* . The computations for each patch are basically the same as described above, except the estimates are expressed as the slant 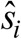 and distance of the patch 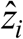, rather than the slant 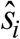 and intercept distance 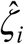 (see Figure 5). Directly using Equation 2 for *i*^th^ image patch gives

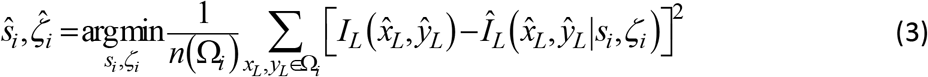

**Figure 5.**
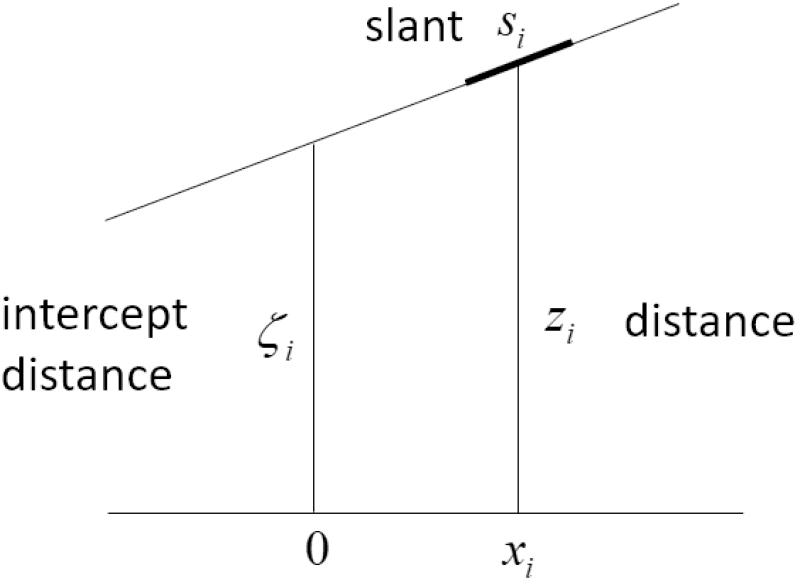
Local planar cross correlation. Illustration of the difference between intercept distance and distance. Local planar cross correlation uses estimates of local slant and distance. Standard cross correlation uses estimates of local distance assuming the local slant is zero.

The problem here is that the estimates of 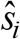 and 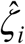 become more correlated the further the horizontal distance of the patch, *x*_*i*_ , from cyclopean axis. On the other hand, 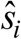 and 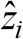 are nearly statistically independent everywhere. The relationship between the intercept distance and distance of the patch is given by *ζ*_*i*_ = *z*_*i*_ − *x*_*i*_ tan*S*_*i*_ . Substituting into the equations in Appendix Figure A1 gives a formula equivalent to Equation 3, but with minimization taken over local slant and distance:

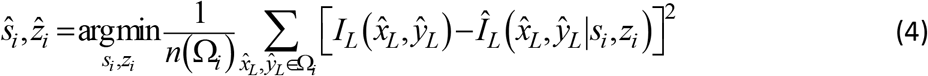

Because the local slant and distance estimates are relatively independent, it is then possible to obtain two independent estimates of the global slant, one from the local distance estimates,

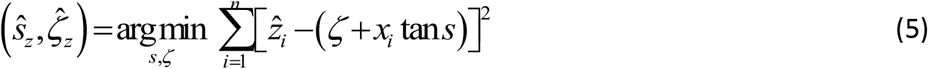

and one from the local slant estimates,

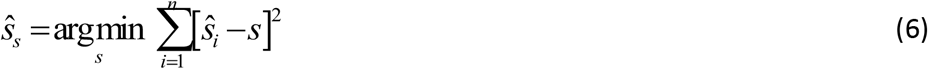

Finally, these two estimates can be combined using reliability-weighted cue combination, which is optimal for bias-corrected statistically-independent cues (Cochran 1937; Clark & Yuille 1990; Oruc et al. 2003):

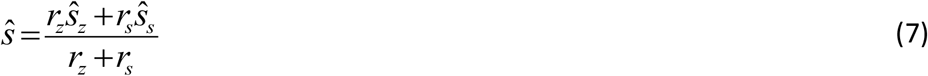

where *r*_*z*_ and *r*_*s*_ are the reliabilities of the two estimates. To measure the reliability of the slant estimates for each cue, we computed slant estimates (for many test patches) separately for each reference slant. From these estimates we then computed the bias in the slant estimates for each reference slant. Finally, the reliability of the estimates was determined from the standard deviation of a cumulative Gaussian fit to bias-corrected slant estimates, as a function of test-patch slant. The reliability of the slant estimates was taken to be one over the square of this standard deviation (i.e. the reciprocal variance).

### Standard Cross Correlation (SCC)

As mentioned earlier, the standard cross-correlation model is a special case of the local planar cross-correlation model where the local slant is assumed to be zero; i.e., Equation 4 with *S*_*i*_ =0:

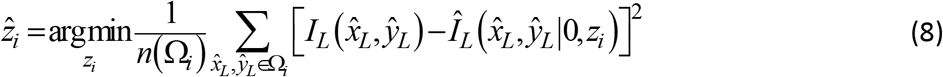

which is equivalent to horizontally translating each patch in the right eye to find the best match in the left eye to obtain an estimated disparity, and then computing the estimated distance given the separation between the eyes (see Figure 2). The estimated slant of the test plane is then obtained by applying Equation 5 to the set of estimated distances.

### Model Predictions

Predictions were generated for two families of model observer. The first family was the models as described above, with no free parameters, except for the patch width *w* used in the LPCC and SCC models (in the PCC model the patch width is the whole test plane). The predicted slant-discrimination thresholds were obtained by simulating model-observer slant estimates as a function of test-plane slant for the four reference slants and three uncorrelated noise levels in the human experiments. On each trial, a model observer returns an estimate of the slant of the test plane. This estimated slant is compared to the fixed reference slant (which was assumed to be known because of the reference surface’s large size, rectangular 3D shape, and absence of uncorrelated noise). If the estimated slant is greater than the reference slant the model observer responds “more slanted.” The proportions of “more slanted” responses were then analyzed to obtain predicted thresholds using the same procedure used for the human responses.

The second family included the same models as before, but two additional free parameters, (i) a fixed level of internal noise *σ*_0_ added to the slant estimates of a model, and (ii) an overall efficiency scale factor *η* that scales all discriminability (*d′*) values of a model down so that the thresholds best match the human thresholds. To obtain predicted thresholds, slant estimates were generated for the specific test plane slants presented to the human observers. For each condition, we computed the mean and the standard deviation of the slant estimates of the model being fitted. From these means and standard deviations, we computed the error rates predicted given any assumed values of the internal noise and scale parameters. The parameters were estimated by maximizing likelihood. The predicted thresholds are calculated from fitted error rates. An equivalent procedure was used for estimating the predicted depth-discrimination thresholds. For more details about the model predictions see https://github.com/CanOluk/Stereo-Slant-Discrimination.

## Results

### Slant Discrimination

Figure 6A shows the scaled slant discrimination thresholds of the three observers (dotted curves) and their average thresholds (solid curves). As can be seen, thresholds increased with the noise contrast (plot color) and decreased with the absolute reference slant (increasing along the abscissa). As noted earlier, the individual human observers exhibited similar patterns of slant-discrimination thresholds and biases, but the thresholds differed substantially in overall sensitivity. To better compare the threshold and bias patterns we separately scaled the thresholds of the most sensitive and the least sensitive observer by an overall factor to best match (in squared error) the thresholds of the medium-sensitive observer; the biases were scaled by the same factors (see Methods).

**Figure 6.**
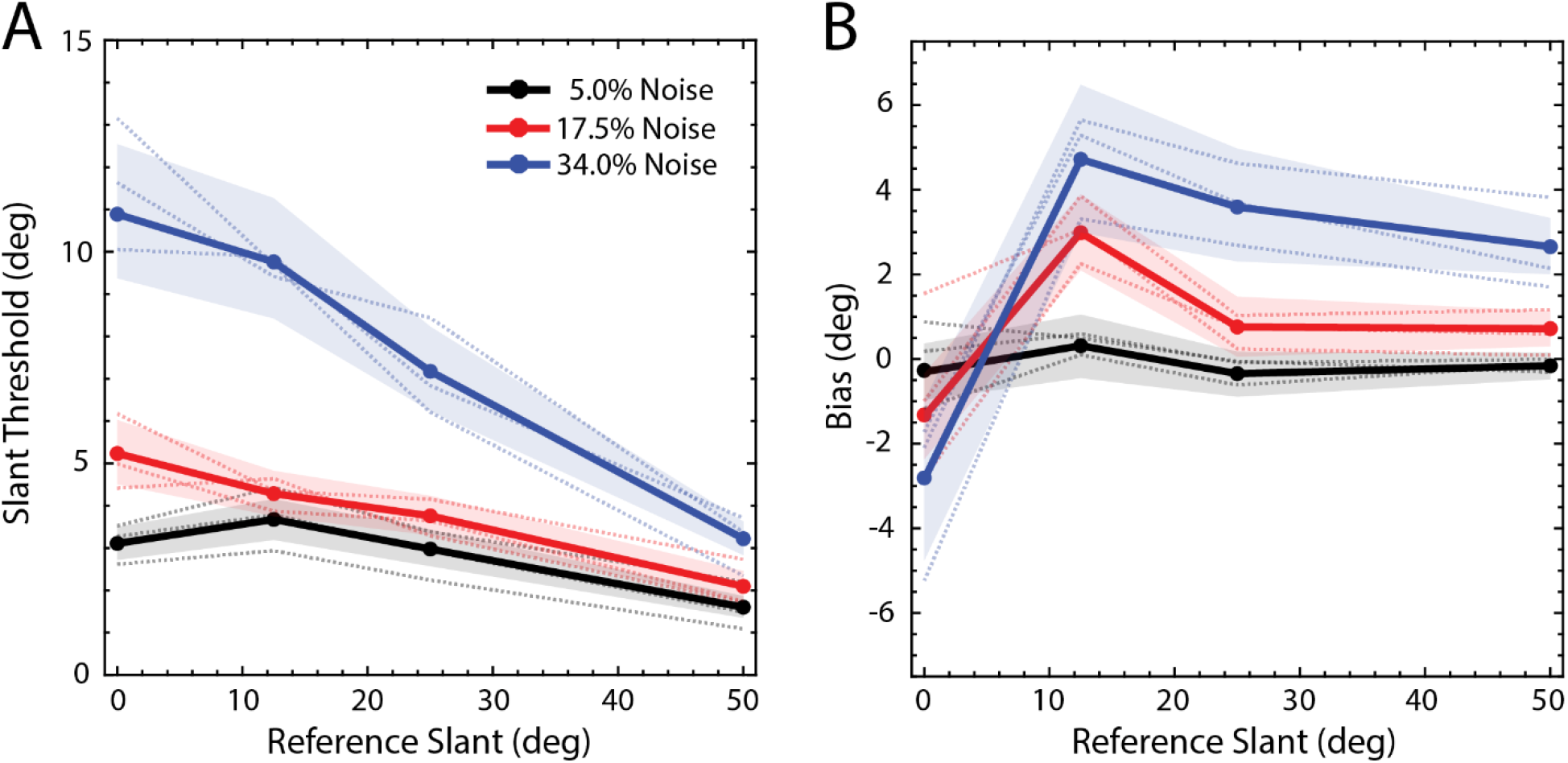
The results of the slant experiment. **A**. Mean thresholds are shown with solid lines and individual thresholds are shown with dotted lines. Ninety-five confidence intervals are shown as shaded regions (see methods for details). **B**. Mean biases are shown with solid lines and individual’s biases are shown with dotted lines. Ninety-five confidence intervals are shown as shaded regions (see methods for details). The individual subjects have similar shaped threshold curves, but they differ in overall sensitivity. In this plot, the most sensitive and the least sensitive observer’s thresholds were each scaled by a single factor to best match the medium sensitive observer’s thresholds. The biases were scaled by the same factors. The two scale factors are 0.68 and 2.01.

Figure 6B shows the biases. When the RMS contrast of the noise was 34% (blue), the frontoparallel test plane (slant of 0°) was perceived as if its right edge was slightly behind the left edge (95% confidence interval: −5.33 to −1.04 degrees), and the slanted test lanes were perceived as slightly more frontoparallel than their true slant (95% confidence intervals of 3° - 6.6°, 2.3° - 5.1°, and 1.9° - 3.3° degrees for reference slants of 12.5°, 25° and 50°, respectively). Similar, but weaker, frontoparallel biases were found for 12.5° and 50° reference slants when the RMS contrast of the noise was 17.5% (95% confidence intervals of 2.2° - 4° , 0.2° - 1°, respectively). A potential explanation for the frontoparallel bias is that the visual system exploits knowledge of the prior probability distribution of slants in natural scenes, which have been found peak at a slant of zero (Burge et al., 2016). When the uncorrelated noise levels are high, the slant estimates necessarily become less reliable. When this occurs, it is rational for the visual system to place more weight on the prior probability distribution, causing a greater bias toward zero slant (e.g., see Weiss, et al. 2002).

Figure 7 shows the absolute thresholds for the first family of model observers along with the most sensitive human observer on a log y-axis. The thresholds of the three model observers are given by the colored symbols. The PCC model has no free parameters. The LPCC and SCC models each had a single free parameter: the patch width *w* . The value of this parameter was set so as to maximize the performance of the SCC model. The value of the patch-width parameter in the LPCC model was identical to the patch width in the best-performing SCC observer. All three models outperform the best-performing human participant in the experiment (black symbols). As expected, the thresholds of the PCC observer (blue symbols) are lowest in all conditions. The performance of the LPCC model is similar to the SCC model; however, if the patch width used for the LPCC observer is made larger, its performance improves, and (of course) asymptotes to the performance of PCC observer.

**Figure 7.**
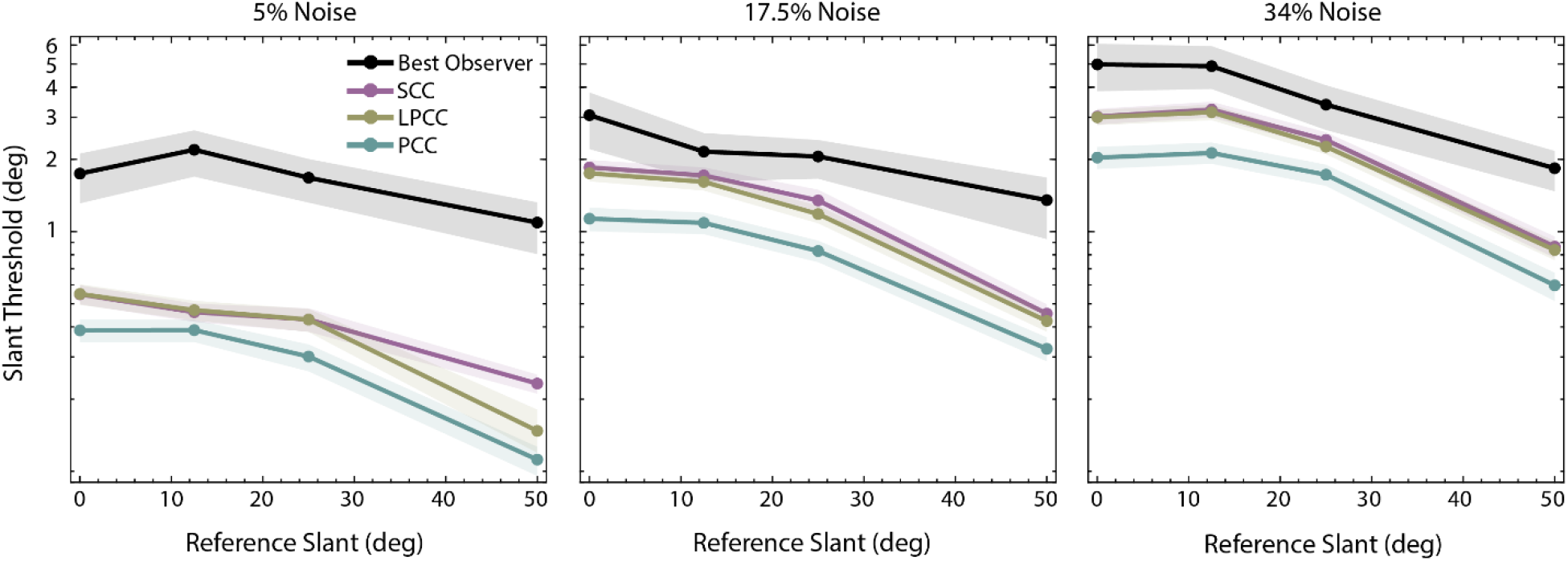
The absolute slant discrimination thresholds of three model observers and the best-performing human observer. The planar cross correlation (PCC) model is near optimal and has no free parameters. The standard cross-correlation (SCC) is suboptimal. The only parameter is the cross-correlation patch width (*w*), which was picked to give the best overall absolute performance (*w* = 0.5°). The local planar cross correlation (LPCC) model is also suboptimal. Shown here is the performance with the same patch width as the best performing SCC model (*w* = 0.5°). Shaded regions correspond to 95% bootstrapped confidence intervals. Note that in this figure (unlike Figure 6) the thresholds are plotted on a logarithmic scale.

Figure 8 shows the thresholds of LPCC and SCC models for different patch width values when the reference slant is either 12.5° or 50° (solid and dashed lines). As patch width increases, the SCC model thresholds become systematically worse than those of the LPCC and PCC models, whereas LPCC model thresholds either improve slightly or remain stable. The LPCC model is therefore more robust to changes in patch width. As expected, the SCC model performs particularly poorly when the reference slant is high because it is the condition when the implicit assumption of frontoparallel surfaces is most inaccurate. Lastly, the difference between LPCC and SCC is highest when the noise is low, probably because the effect of external noise tends to exceed the effect of the differences in specific computations.

**Figure 8.**
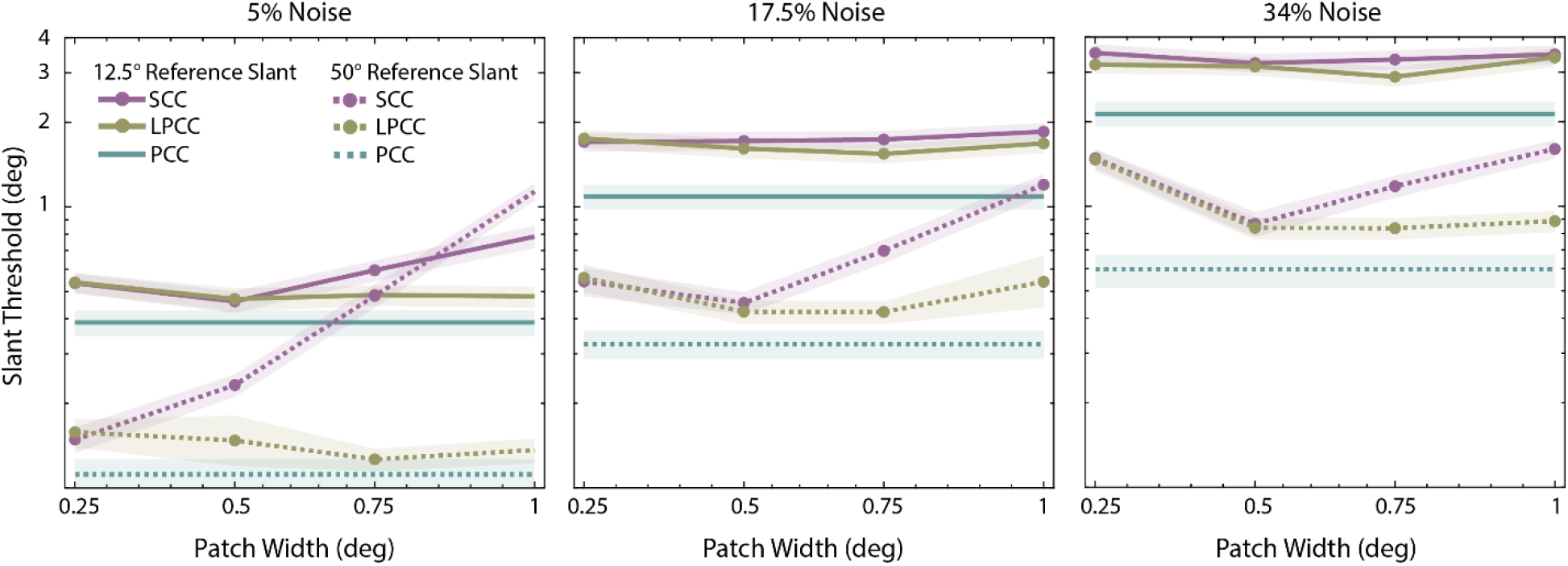
The change in absolute slant discrimination thresholds of LPCC and SCC model observers as function of patch width. The patch width for the PCC model was the full size of the target in the right eye (2.2 °). The solid lines correspond to 12.5 ° reference slant and dashed lines correspond to 50 ° reference slant. Shaded regions correspond to 95% bootstrapped confidence intervals. Note that the thresholds are plotted on a logarithmic scale.

Figure 9 shows the maximum-likelihood fits of the three models (dashed curves) to the average thresholds in Figure 6A, when the patch width (*w*), estimation-noise standard deviation (*σ*_0_) and overall efficiency scale factor (*η*) were allowed to vary (only *σ*_0_ and *η* were allowed to vary for the PCC model). Note that although the fits were obtained by maximizing likelihood, we report the root-mean-squared error (RMSE) in the figure because it is more intuitive. The predicted rate in fall-off in the thresholds with reference slant is similar to that in the human observers. Note that the best fitting values of *σ*_0_ are quite small, on the order of 1° − 1.5°, and hence are in a plausible range. Surprisingly, the predictions are about equally good for the three models, suggesting that the data are not sufficient to differentiate between the models.

**Figure 9.**
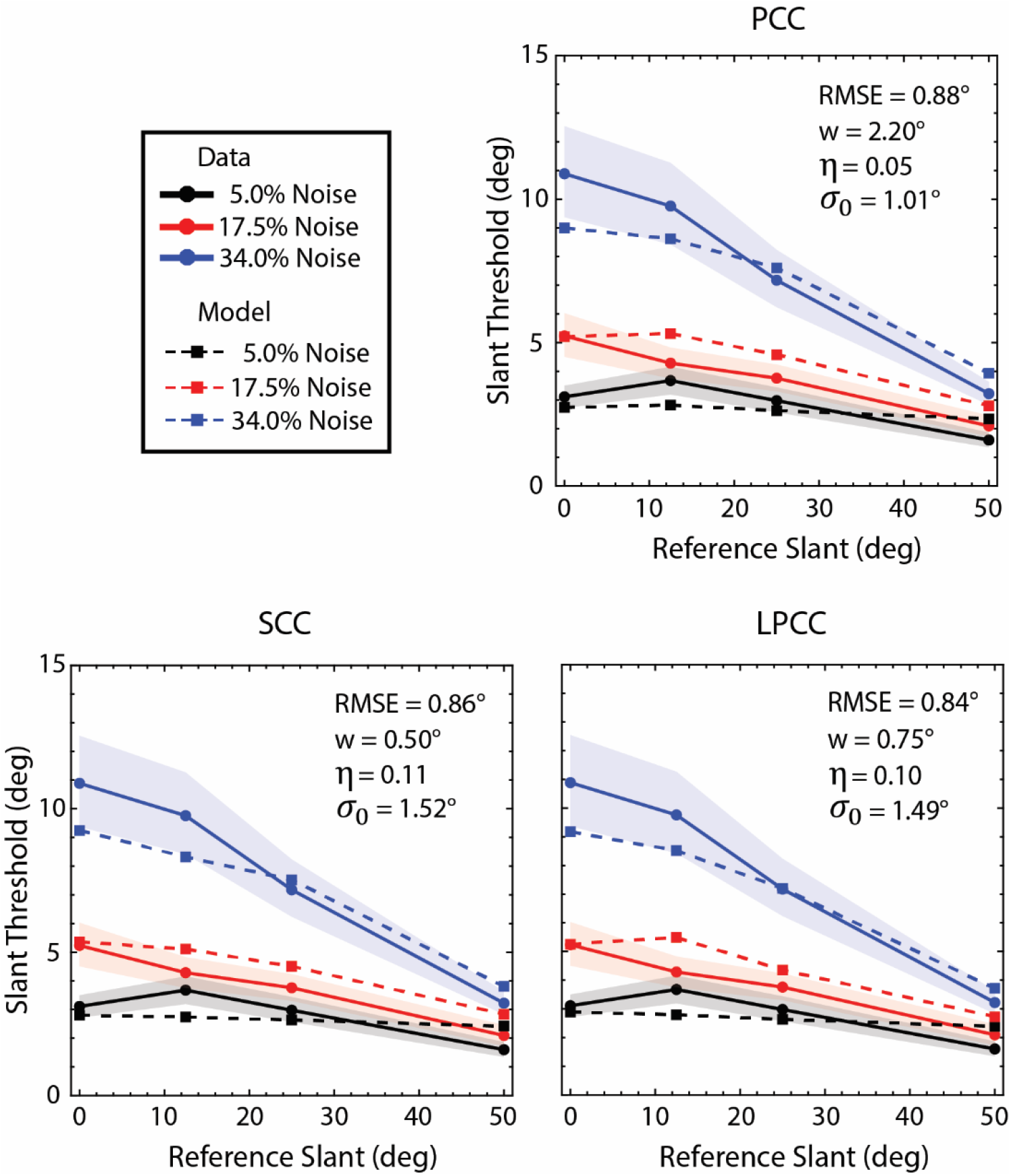
Model fits. For each model observer, the patch width (*w*), scale factor (*η*), and estimation noise standard deviation (*σ*_0_) were estimated by maximizing the likelihood of the data. The patch width for the PCC model was the full size of the target in the right eye (2.2°). Although the parameters were estimated by maximizing likelihood, the goodness-of-fit measure shown in the figure is the RMSE, which is more intuitive.

Figures 10A and 10B plot goodness-of-fit measures (RMSE and negative log-likelihood respectively) as a function of patch width. The arrows in Figure 10A indicate the patch widths of the predictions shown in Figure 9. They correspond to the best fits in terms of RMSE. Both plots show that including the estimation-noise parameter greatly improves model predictions (shaded regions are 68% confidence intervals). Figure 10C shows the maximum-likelihood-parameter estimates of each model (symbol color) for each patch width (symbol size). The estimated noise and scalar parameters are largest for the SCC model, smaller for the LPCC model, and smallest for the PCC model.

**Figure 10.**
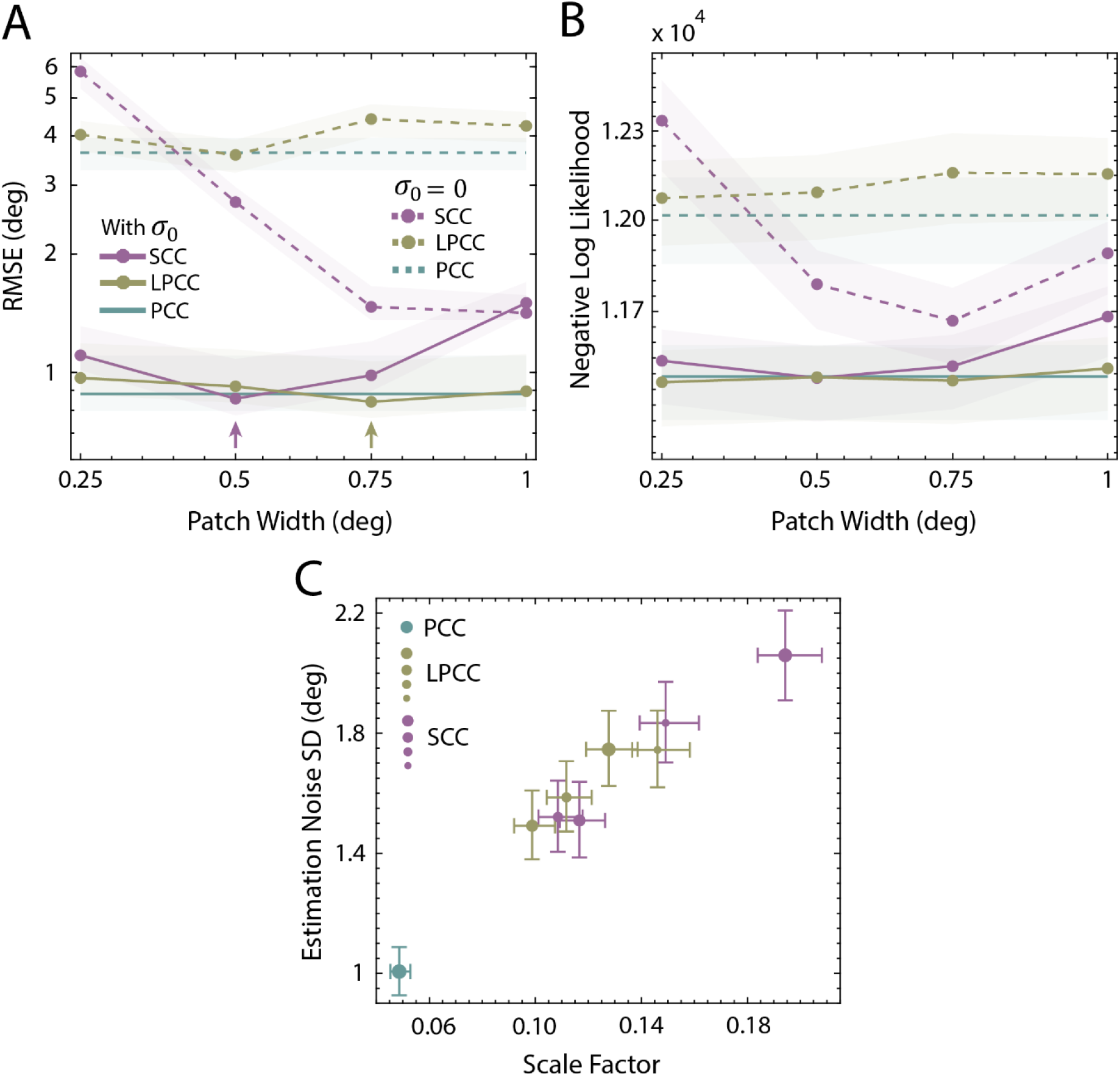
Results of the model fitting procedure. **A**. Root-mean squared error (RMSE) for the three models as a function of patch width. The arrows indicate the smallest RMSE for the SCC and LPCC models. The parameters were actually estimated by maximizing likelihood (minimizing negative log likelihood). **B**. Negative log likelihood for the three models as a function of patch width. For the SCC model the RMSE and negative log likelihood are minimal at the same patch width. For the LPCC model the RMSE is minimal at 0.75 but the negative log likelihood is minimal at 0.25. **C**. Estimated scale factor and estimation-noise standard deviation for the three models. For the LPCC and SCC models, increasing symbol size represents increasing patch width (0.25, 0.5, 0.75, and 1 visual degree respectively). Error bars are 68% bootstrapped confidence intervals.

In a control experiment, participants performed the slant discrimination experiment with the right eye covered to determine the usefulness of monocular cues. The tested slant was 50° because this was the largest slant magnitude in the main experiment and because perspective (monocular) cues are strongest in this case (Knill, 1998). Psychometric functions were measured for all three noise levels. It was not possible to estimate thresholds, because the slopes of the psychometric functions were always near zero. We conclude that the human slant discrimination thresholds were based entirely on binocular (stereo) cues.

### Depth Discrimination

Figure 11A shows the depth discrimination thresholds of the three observers (dotted curves) and their average (solid curve), and Figure 11B shows the biases. The individual differences were less in this experiment than in the slant-discrimination experiment, and hence there is no scaling of the thresholds for the least and most sensitive participants. As can be seen, human thresholds increase with noise contrast. When the noise is highest (34% RMS), there is also a small bias in two of the subjects to see the test plane slightly farther than the reference plane (the 95% confidence interval of the average bias is 3.9 to 19.8 arc sec).

**Figure 11.**
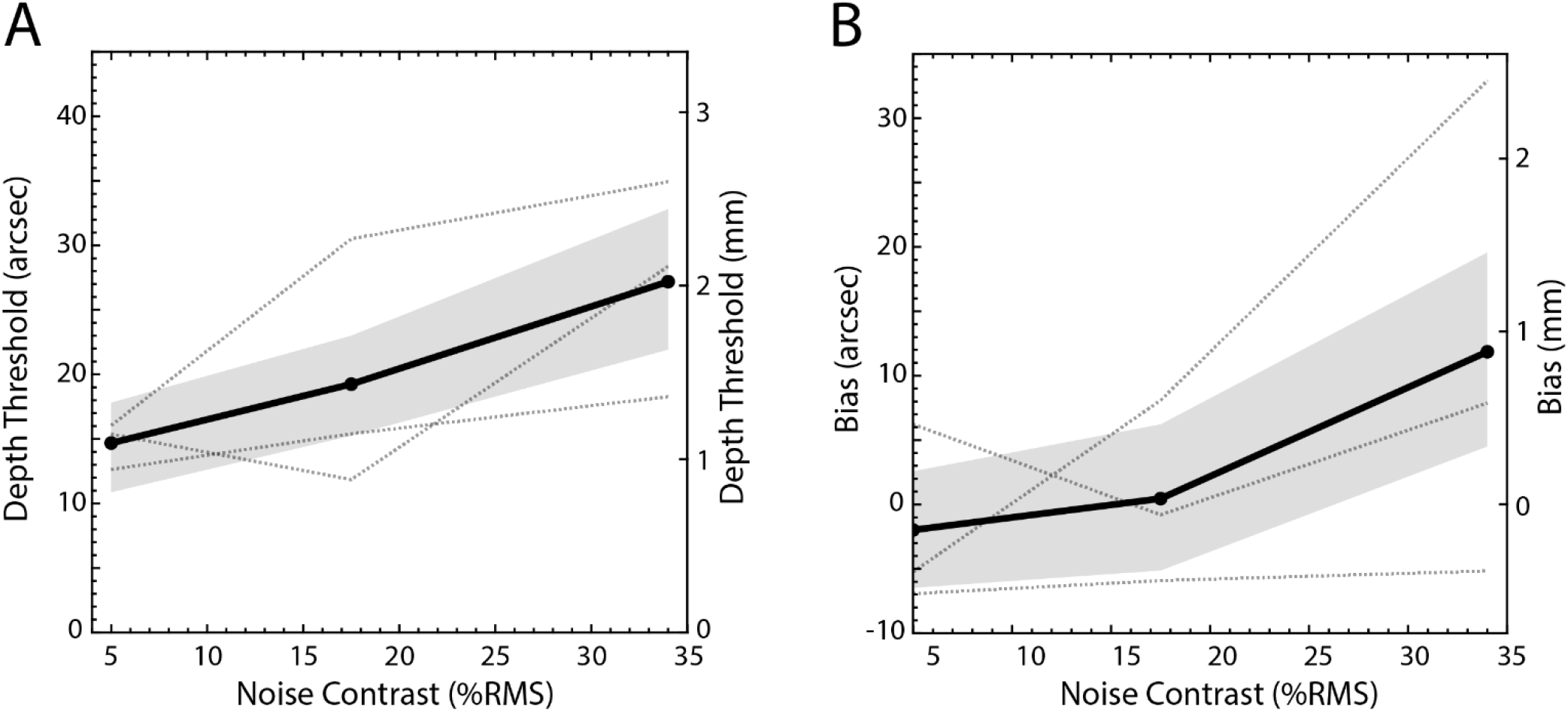
The results of the depth experiment. **A**. Mean thresholds are shown with solid lines and individual thresholds are shown with dotted lines. Ninety-five confidence intervals are shown as shaded regions (see methods for details). **B**. Mean biases are shown with solid lines and individual’s biases are shown with dotted lines. Ninety-five confidence intervals are shown as shaded regions (see methods for details).

Similar to the slant experiment, the depth discrimination thresholds of the optimal PCC model are better than those of the LPCC and SCC models (1.3 seconds of arc better on average). Thresholds of LPCC and SCC models are generally similar but the performance of LPCC is considerably better than SCC in the low noise condition for 0.75 ° and 1 ° patch widths (0.5 and 1.5 seconds of arc better respectively).

The maximum likelihood fits of the three models to the data revealed similar results to the slant experiment. Without the estimation noise parameter, the three models do not fit the thresholds well (RMSE: around 30 seconds of arc, or 2.2 mm). When the estimation noise parameter had a standard deviation 5-7 seconds of arc (0.3-0.5 mm), all three observer models fit the thresholds quite well (RMSE: approximately 1 second of arc or 0.07 mm).

Recall that in the depth experiment the slant of the reference plane was zero, but the test plane was slanted at 44°. This was done so that the pedestal-disparity values were similar to those in the main experiment, making it easier to compare human efficiency for slant and depth discrimination. The PCC observer is approximately ideal for both slant and depth discrimination. Thus, it is possible to compare absolute efficiencies in the two tasks. The efficiency scale factors for aligning the PCC thresholds with each participant’s thresholds in the high noise conditions are shown in Figure 12A. For two participants (P1 and P2), there is a trend for human efficiency to be higher in the slant discrimination experiment.

**Figure 12.**
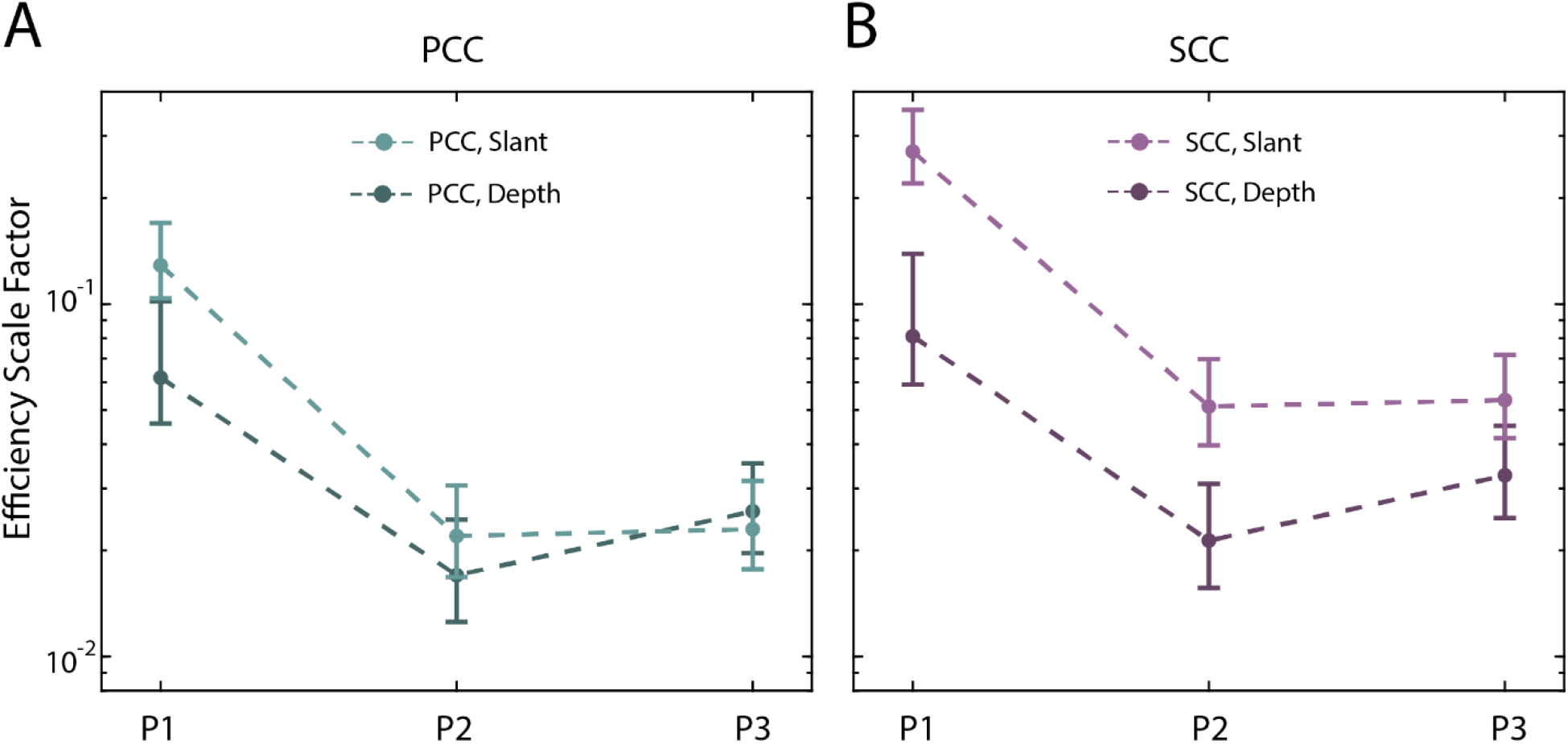
The efficiency scale factors estimated from two different experiments. **A**. Using the PCC Model, for three participants, scale factors estimated by maximizing the likelihood of the data of the highest noise condition either for slant (50-degree reference slant condition) or depth experiment. 68% bootstrapped confidence intervals are shown with error bars. **B**. Using the SCC model observer, for three participants, scale factors estimated by maximizing the likelihood of the data of the highest noise condition either for slant (50-degree reference slant condition) or depth experiment. 68% bootstrapped confidence intervals are shown with error bars.

It is also possible to estimate the efficiency scale factors for the SCC model. For all participants, the efficiency is higher in the slant experiment, and the confidence intervals do not overlap (Figure 12B). Overall, humans appear to be more efficient at estimating surface slant than surface distance.

## Discussion

We derived the approximate binocular ideal observer for discrimination of the 3D orientation and distance of textured planar surfaces viewed in the presence of additive white noise that is uncorrelated across the two eyes. This planar-cross-correlation (PCC) observer first filters the left and right images, based on the known frequency content of the texture, to remove irrelevant frequency components due to the white noise. Using projective geometry, the PCC observer then generates a predicted left image from the right image (or vice versa) for each possible 3D orientation and distance of the test plane. The estimated surface orientation and distance are the values that make the most accurate prediction of the left image (i.e., give the smallest mean-squared error). We also considered two sub-optimal observers that also pre-filter the left and right images. The local planar cross correlation (LPCC) observer uses the same fundamental computations as the PCC observer, but it makes multiple local estimates of the 3D orientation and distance in local image regions, and then combines those local estimates to obtain a single estimate of the 3D orientation and distance of the entire test plane. The standard cross-correlation (SCC) observer uses projective geometry to estimate the distance of local image regions under the assumption that the local surface slant is zero. It then combines those estimates to obtain an estimate of the 3D orientation and distance of the test plane. Each of the three model observers has two free parameters: an overall efficiency parameter and a parameter representing a fixed level of internal estimation noise.

In terms of absolute performance, the PCC observer performs substantially better than the LPCC and SCC observers, because it optimally combines all the disparity information over the approximately 2° × 2° test plane. The LPCC observer performs better than the SCC observer, especially for larger patch sizes and slants, and it is also less affected by patch size and surface slant (it is more robust) than the SCC observer.

We compared the performance of these model observers with human observers in a stereo slant-discrimination experiment. Human thresholds were measured as a function of the reference slant and the noise contrast of separate samples of white noise added to the left-eye and right-eye images. Human thresholds decreased with the slant of the reference plane and increased with the level of uncorrelated noise. The pattern of thresholds was consistent across the three human observers. In control experiments, we found that the human thresholds were based on stereo cues alone, and that there is a trend for humans to be more efficient at slant discrimination than depth discrimination. All three models were able to predict the measured thresholds with approximately equal quantitative accuracy.

### Binocular differences and the correspondence problem

One way to describe the difference between left and right images (for the imaging geometry in Figure 2) is in terms of corresponding points: for planar (and many other) surfaces, each point on the surface projects to corresponding points in the right and left images. The corresponding image points can be described by a single horizontal disparity (translation). If one knows all of the corresponding points (i.e., all the horizontal disparities), as well as the geometry of the imaging surfaces for the two eyes, then one has a complete description of the binocular information concerning the orientation and distance of the planar surface. Thus, one hypothesis for how the visual system estimates surface orientation and distance is that it first determines all the horizontal disparities (corresponding points) and then estimates the surface orientation and distance from the set of disparities.

Determining the corresponding points can appear difficult if one literally thinks about finding a “point” in one eye that corresponds to a point in the other: the so-called “correspondence problem”. Various proposals for how to solve the correspondence problem have been tested in the human and computer-vision/image-processing literature. In most, some region around each point is selected in one eye’s image and then translated (and perhaps also distorted) to find the best matching region in the other eye’s image. The horizontal translation between the centers of the best-matching regions in the two eyes is the estimate of the horizontal disparity (the corresponding point) for that point. Ironically, random element stereograms of frontoparallel surfaces, once considered the ultimate demonstration of the correspondence problem, are trivial to match using such local region translation methods (i.e., standard cross correlation).

Another way to describe the differences between the left and right images is by describing the back-projection mapping between the spatial pattern of gray levels in the two images. Planar cross correlation (PCC) obtains this description by finding the 3D surface orientation and distance that best explains the difference between the two images. The estimated horizontal disparities of the corresponding points are implicit in the estimated surface orientation and distance, but are never directly estimated. The same logic holds for local planar cross correlation (LPCC). It finds the 3D orientation and distance that best explains the difference between image patches in the two images. Although standard cross correlation (SCC) can be described as a special case of local planar cross correlation, it is most natural to conceptualize it as the traditional approach of first directly solving the correspondence problem (estimating the disparity) for each image point and then estimating the surface distance and orientation from the disparities.

Both ways of describing the differences between the left and right images are valid, but they suggest different hypotheses for neural mechanisms. The first description most naturally leads to the hypothesis that early receptive fields are explicitly coding binocular differences in horizontal phase and position (horizontal disparities), and that these are integrated in later areas to yield receptive fields sensitive to distance and surface orientation.

The second description most naturally leads to the hypothesis that early binocular receptive fields are explicitly coding the spatially structured patterns of binocular differences that are produced by back-projection of planar surfaces. For example, Figure 13 shows the binocular receptive fields that would respond best to a sinewave textured surface at 100 cm, with a slant of 45 deg, for five different tilts. To emphasize the shape differences between to left and right receptive fields, the imaging plane was set to the same distance as the surface (100 cm). If the imaging plane was instead set at plus or minus 17 mm (the location of the retinas), then the left and right receptive fields would also differ in position. When the surface tilt is 0 deg (see Fig. 1B), the left and right receptive fields differ primarily in scale/frequency. When the surface tilt is 90 deg the left and right receptive fields differ primarily in orientation. For other tilts there are scale, orientation and shear differences (see also Figure 3). The third column shows the differences between the left and right receptive fields. Figure 12B shows how the total energy of the difference of the left and right receptive fields varies with slant and tilt. The energy tends to increase with slant and decrease with tilt, although this latter effect is quite small for low slants.

**Figure 13.**
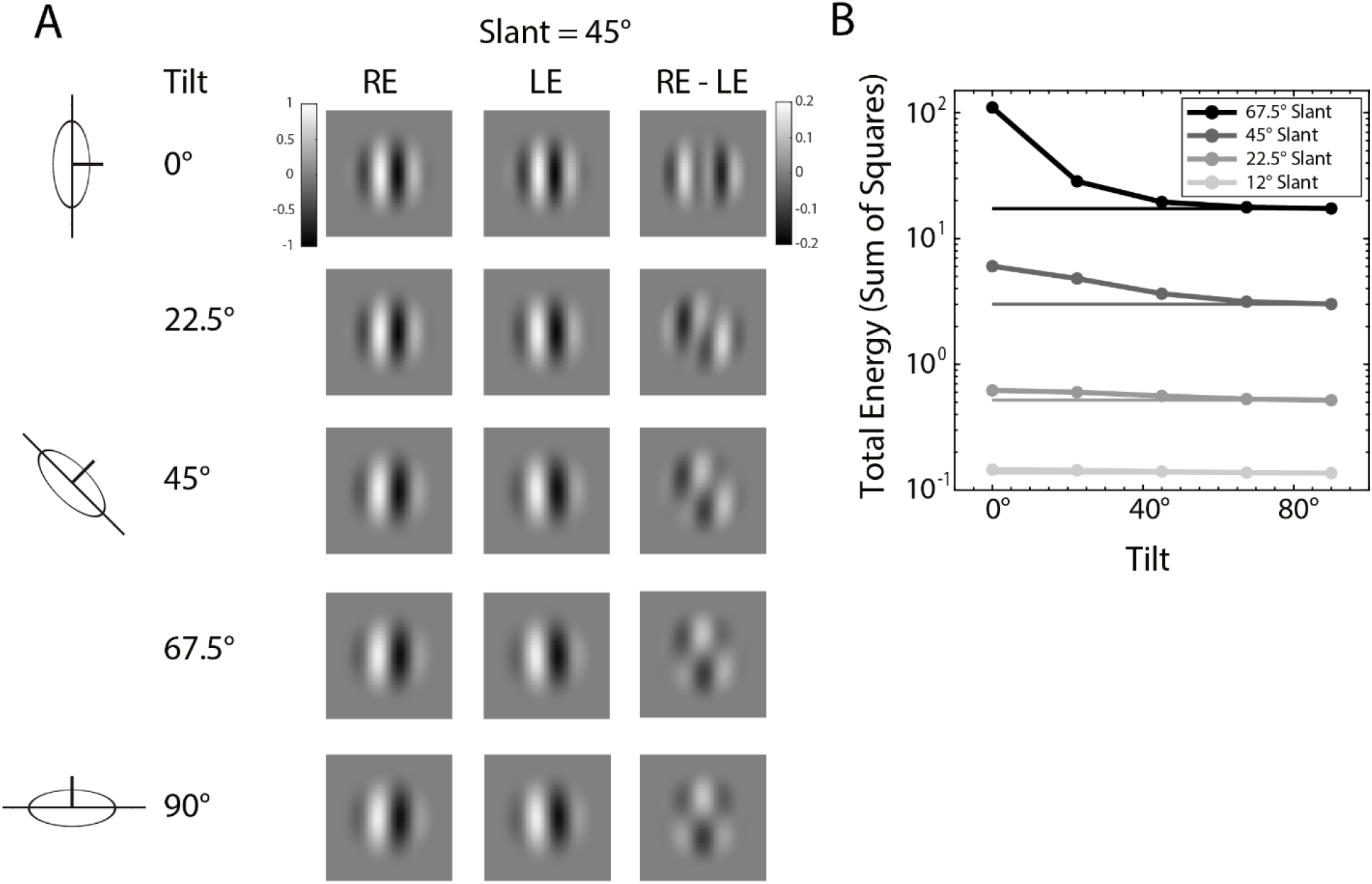
Hypothetical binocular receptive fields tuned to surface orientation and distance. **A**. Examples: right eye receptive field (RE), left eye receptive field (LE) , difference between right and left eye receptive fields. **B**. Energy of the difference between right and left receptive fields as a function of tilt, for several different slants.

Although the LPCC and the PCC observers demonstrate that structured disparity patterns provide substantial additional information for slant discrimination and estimation (see also, Jones & Malik, 1992; Super & Klarquist, 1997; Ogale & Aloimonos, 2005; Li & Zucker, 2010; Vidal-Naquet & Gepshtein, 2012), there remains uncertainty about the extent to which structured disparity patterns are explicitly exploited in the visual system, and at what stages of visual processing.

Single unit recordings in primary visual cortex of monkey and cat have found populations of neurons with binocular receptive fields that are consistent with structured disparity patterns in orientation (Bridge & Cumming, 2001) and spatial frequency (Sanada & Ohzawa, 2006). However, modeling with generalized versions of the disparity energy model originally introduced by Ohzawa et al. (1990) showed that most of the useful disparity information is carried by standard horizontal disparity detectors (Bridge & Cumming, 2001; Bridge et al., 2001; Sanada & Ohzawa, 2006). Nonetheless, these models do not consider all of the structured disparity patterns associated with planar surfaces (see Figure 13). Also, the possible benefits of including information about structural disparity patterns may better emerge in models of population decoding that pool efficiently over all the relevant neurons (Bridge & Cumming, 2008; Greenwald & Knill, 2009; Kato et al., 2016).

Psychophysical studies have also not yet provided a clear picture of the role of structured disparity patterns. Studies that have focused on structured disparity patterns have revealed that it is difficult to reject simple SCC type models (Halpern et al., 1996; Hibbard & Langley, 1998; Greenwald & Knill, 2009). Studies that have either reported evidence for SCC-type computations (Banks et al., 2004; Filippini & Banks, 2009; Allenmark & Read, 2011; Goutcher & Hibbard, 2014) or for LPCC-type computations (Hibbard et al., 2002), generally have not directly compared the models. Thus, there does not appear to be compelling evidence either for or against the visual system’s use of structured disparity patterns. Nonetheless, we (and others) have shown that PCC and LPCC type computations are more accurate at slant estimation under naturalistic conditions than the simple SCC computation. Thus, there should have been evolutionary pressure to incorporate similar computations into the early visual system.

Finally, it is important to note that while the PCC model is approximately optimal for estimating the slant and distance of planar textured surfaces (with uncorrelated image noise), it is not optimal under real-world conditions, where many surfaces are non-planar and where there are half occlusions (points with no corresponding point in the other eye). More sophisticated computations are required in natural 3D scenes (Scharstein & Szeliski, 2002; Hirschmuller & Scharstein, 2007). The human visual system is likely to be much more sophisticated than the models considered here.

There are additional research approaches that could be useful for discriminating between the different model described here. One approach is to vary stimulus parameters, such as the size and spatial-frequency content of the test patches and the range of test slants and tilts, to find those parameter values where the models make the biggest differences in the predicted pattern of slant thresholds. Testing with these parameter values should better differentiate between the models. Another approach is to consider what specific computations are optimal for natural images. For example, Burge & Geisler (2014) used accuracy maximization analysis (AMA) of natural images to determine the set of vertically-oriented binocular receptive fields that are optimal as a population for estimation of local disparity. The receptive fields in this population share many properties with receptive fields measured in visual cortex. It should be possible to perform a similar analysis to determine the optimal population of binocular receptive fields for estimating surface slant or surface slant and distance of natural images. This type of normative analysis could be used to guide investigations of receptive-field properties in cortex that may capitalize upon the structured disparity patterns that result from binocularly-viewed slanted surfaces.

### Coordinate systems and the representation of surface orientation

The equations in Appendix Figure A1 are based on projecting the scene onto an image plane. This is the common framework for representing camera images and leads to the simple equations used here. In vision science, it is also common to represent images in spherical coordinates, which is equivalent to projecting the scene onto spherical surfaces centered on the nodal point of each eye. If desired, it is straightforward to convert the equations in Figure A1 into spherical coordinates (azimuth and elevation) by substitution. It is also common in vision science to consider cases where the eyes are not in primary position (pointing straight ahead). In the Appendix we provide formulas for generalizing the closed-form expressions in Figure A1 to the case where each eye is rotated arbitrarily about the three axes.

Throughout this manuscript, estimates of surface orientation and distance were made in a headcentric coordinate system. The matching computations could also be used to make estimates in direction-centered coordinate systems. For example, the direction-centered coordinate system for each image location could be defined as one aligned with the axis passing through that image location and the origin—direction-centered coordinate systems are illustrated in Figures 2C. A plausible hypothesis is that local surface orientation and distance are initially estimated in local direction-centric coordinates. These estimates might be used when the observer is tasked with reporting local surface slants with respect to a direction. These local measurements might then be mapped into headcentric coordinates, where they are grouped into 3D surfaces. Importantly, the grouping rules are often simpler in headcentric coordinates than in direction-centric coordinates. For example, with the representation in Figure 2B, one can use simple similarity grouping to group estimated local slants into a 3D plane, but similarity grouping would not work for the representation in Figure 2C. Also, as Backus et al. (1999) note, mapping into head-centric and body-centric representations are important for implementing motor behaviors.

## Conclusion

Stereo slant discrimination performance was measured for accurately-rendered textured surfaces designed so that performance is dominated by binocular-disparity cues. We compared human performance with model observers (PCC and LPCC) that simultaneously estimate distance and surface orientation without directly estimating disparities and with a model observer (SCC) that first directly estimates disparities and then combines those to estimate distance and surface orientation. We found that the pattern of human slant-discrimination thresholds was predicted equally well by all three models. However, we find that the PCC and LPCC models perform substantially better at slant discrimination; hence, there should have been evolutionary pressure to incorporate similar computations into the early visual system. We also mention additional research approaches that may better differentiate between the models.

## Acknowledgements

Supported by NIH grant EY01147 to WSG and JB, NIH grant EY020592 to LKC, and NIH grant EY028571 to JB. Supported by NSF GRFP and Donald D. Harrington Doctoral Fellowship to KB.

## Appendix

## Projection of planar surfaces

Here we derive the exact equations for mapping a point in the right image to the corresponding point in the left image given a planar surface at an intercept distance of *ζ* , with a slant of *s* and a tilt of *τ* , for the imaging geometry illustrated in Figure 2. The equations are shown in Figure A1. These equations are for arbitrary slant, tilt, and distance, but for the current experiments the tilt was set to zero.

Planar cross correlation (PCC) is implemented by applying these equations to all the points in an image patch in the right eye for each possible intercept distance, slant, and tilt of the planar surface. In other words, the predicted image value (e.g., gray level) at the predicted left image location 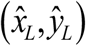 is the image value at 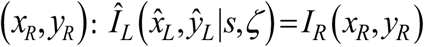. The estimate of distance and surface orientation are the values of distance, slant, and tilt that give the most accurate prediction of the left-eye image (smallest mean squared error).

For the PCC model the image patch is the whole right image of the test plane. For local planar cross correlation (LPCC) and standard cross correlation (SCC), the right image patch is a smaller fixed-size image patch. Also, for the LPCC and SCC models, the equations are expressed in terms of distance *z* rather than intercept distance *ζ* (see Figure 5). The equations in Figure A1 can be expressed in terms of distance by setting *ζ* = *z* − *x*tan*s*cos*τ* + *y*tan*s*sin*τ*. Note that in the present experiment, where tilt is zero, *ζ* = *z* − *x*tan*s* , and that for the SCC model *ζ* = *z* , because of the assumption that *s* =0.

**Figure A1.**
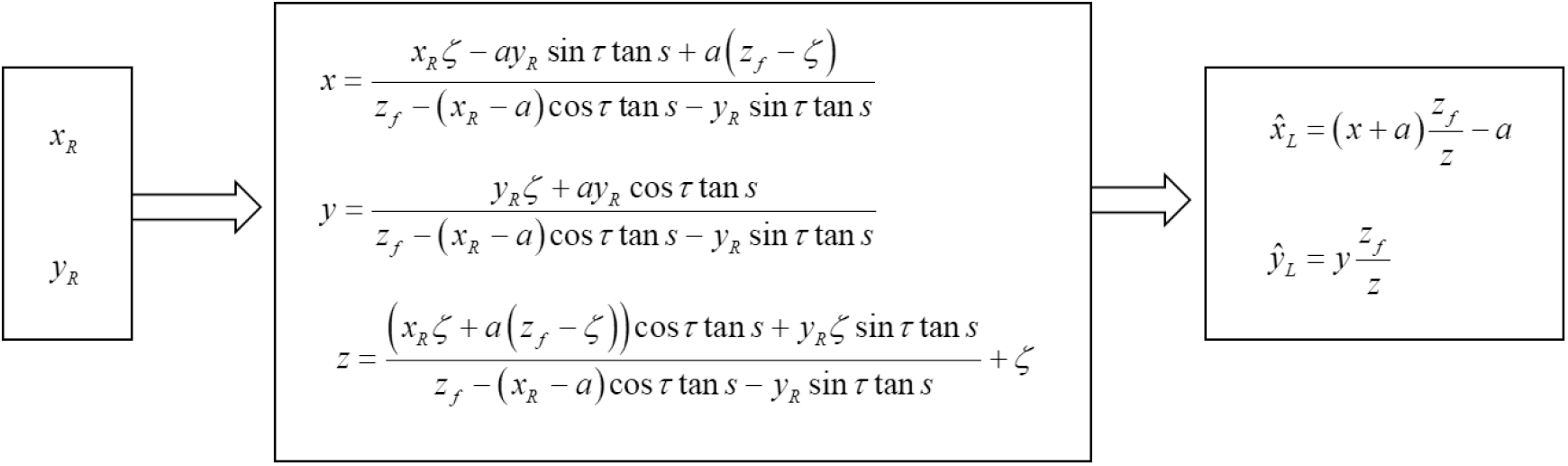
Equations for projecting a point in the right image to a point in the left image given a planar surface at an intercept distance of *ζ* having a slant of *s* and a tilt of *τ*. The distance of the image plane to the nodal point of each eye/camera is *z*_*f*_ and the separation between the nodal points is 2*a* . The predicted image value (e.g., gray level) at left image location 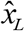, 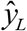 is the image value at *x*_*R*_, *y*_*R*_ .

The equations in Figure A1 are standard projective geometry, but are derived here to provide compact equations that are easy to apply. To derive the equations in Figure A1, we first note that if the nodal point is at the origin in 3D Euclidean space (as in Figure 2), then the standard equations for perspective projection are

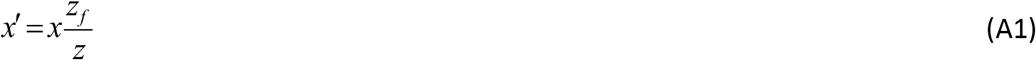

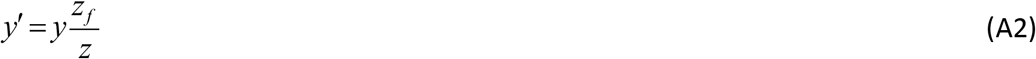

where *z*_*f*_ is the distance from the nodal point to the image plane, and (*x′*, *y′*) is the point in the image plane. If the nodal point is shifted to the left by a distance of *a* (left eye point in Figure 2), then the point in the image plane (*x*_*L*_, *y*_*L*_) is given by

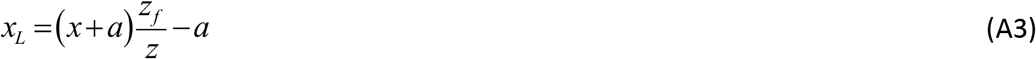

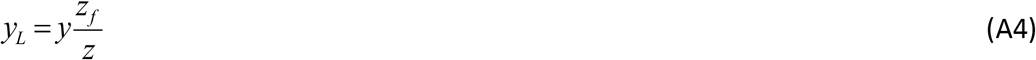

Equations A3 and A4 are the equations in the right panel of Figure A1. If the nodal point is shifted to the right, then the point in the image plane is given by

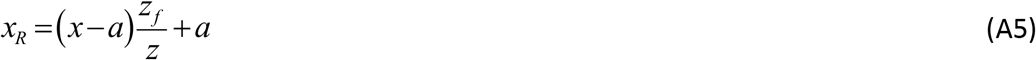

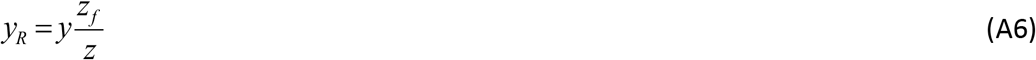

The slant, tilt and distance at a surface point (*x*, *y*,*z*) are defined here in global Euclidean coordinates as the slant, tilt, and intercept distance (*s*,*τ*,*ζ*) of the plane passing through that surface point (Figures 2), where the slant, tilt, and distance are with respect to the cyclopean axis. The intercept distance *ζ* is the intercept of the plane with the cyclopean axis. The slant *s* is defined as the magnitude of the angle (0 – 90°) between the surface normal and the cyclopean (or optic) axis, and the tilt *τ* is defined as the direction (−180° − 180°) around that axis in which distance is changing most rapidly (the counter-clockwise angle of the parallel projection of the surface normal vector; see Figure 1). Using these definitions, the equation of the plane is

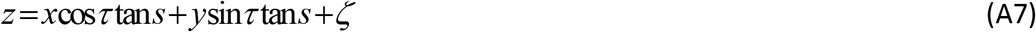

Specifically, the textbook definition of the equation of a plane is **n**□(**x**−**x**_*p*_)=0, where **n** is the normal vector at location **x**_*ζ*_ =(0,0,*ζ*). The normal vector is obtained by rotating the unit normal vector of the frontoparallel plane, (0,0,−1), around the vertical (*y*) axis by angle *s* , and then rotating the resulting vector around the distance (*z*) axis by angle *τ* . Substituting the rotated normal vector into the equation for a plane gives Equation A7.

To derive the equations for back projection, we first rearrange Equations A5 and A6:

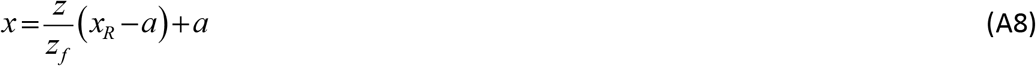

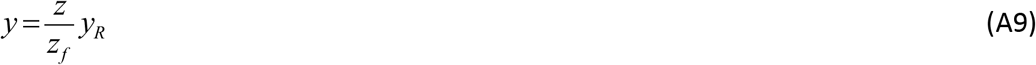

Substituting Equation A7 into Equation A8 we have

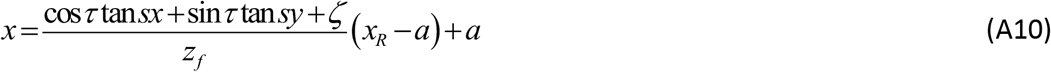

Subtracting *a* from both sides of Equation A8 and then taking the ratio with Equation A9 we get

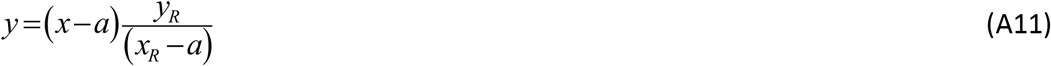

Substituting Equation A11 into Equation A10 gives

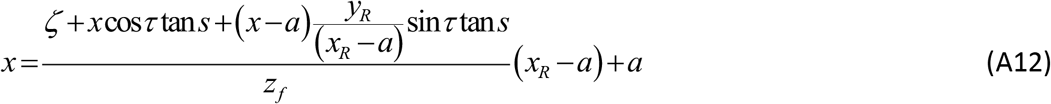

Simplifying this equation gives the expression for *x* in Figure A1,

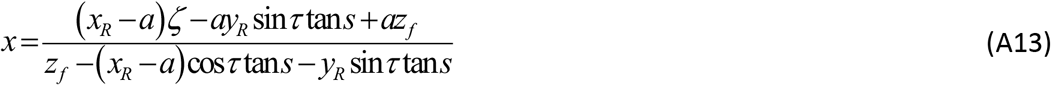

Next substitute Equation A7 into Equation A9

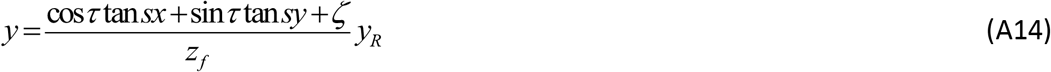

Rearranging Equation A11 gives

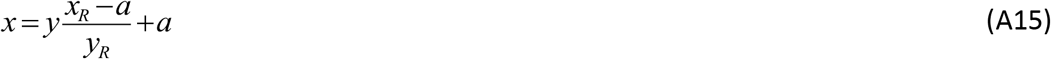

Substituting Equation A15 into Equation A14 and simplifying gives the expression for *y* in Figure A1:

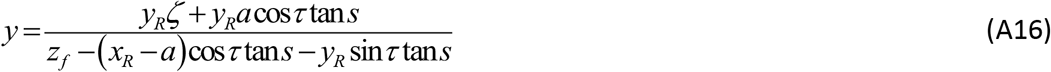

Finally, substituting Equations A13 and A16 into Equation A7 and simplifying gives

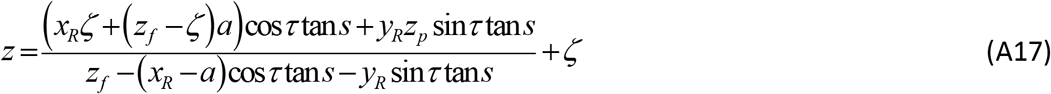

Equations A13, A16 and A17 are the back-projection equations in Figure A1.

## Spherical coordinates

The binocular equations in Figure A1 can be expressed in spherical coordinates (azimuth-longitude *α* and elevation-latitude *e*; Fick coordinates) with respect to the nodal point for each eye by substituting for *x*_*R*_, *y*_*R*_, *x*_*L*_, *y*_*L*_ using the equations:

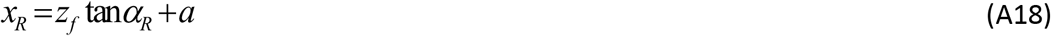

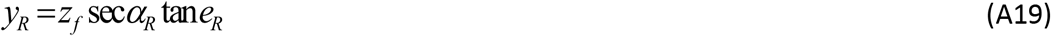

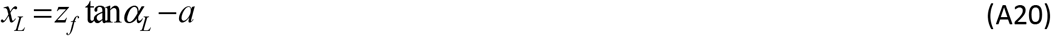

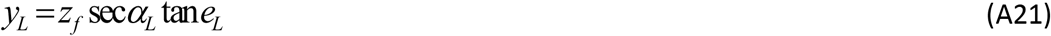

These equations can be derived by starting with the textbook formulas, *x′* = *R*cos*e′*sin*α′* , and *y′* = *R*sin*e′* , where 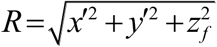 , and then solving for *x′* and *y′* . This “gun-turret system” is a popular spherical coordinate system that is sometimes regarded as best for specifying disparities (Howard & Rogers 2012), but similar equations can be written down for any other spherical coordinate system (e.g., the azimuth-latitude, elevation-longitude system; Helmholtz coordinates).

## Eyes not in primary position

In the human visual system, the eyes can rotate around the three axes (and may even translate by a small amount when rotated). The possible rotations are largely constrained by the modified Listing’s law (e.g., see Howard, 2012), but for the purpose of generalizing the expressions in Figure A1 to different eye positions, we can allow each eye to rotate arbitrarily (Figure 2A). Specifically, suppose that the right eye is rotated from the primary position by angles, *θ*_Rx_ ,*θ*_Ry_ ,*θ*_Rz_ , applied in a specific order, and that the left is rotated by angles, *θ*_Lx_ ,*θ*_Ly_ ,*θ*_Lz_ applied in the same specific order. For example, suppose the order corresponds to the order they are listed above (rotate around x, then around y, and then around z). The rotation matrices for the three directions are given by

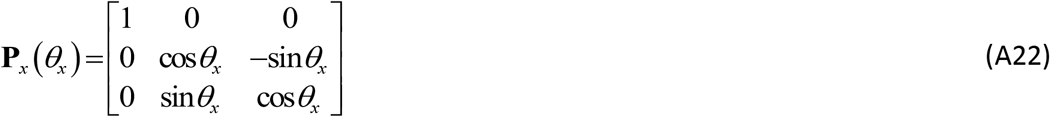

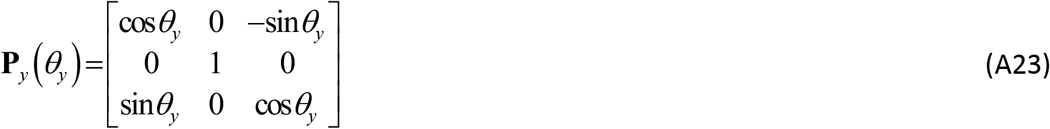

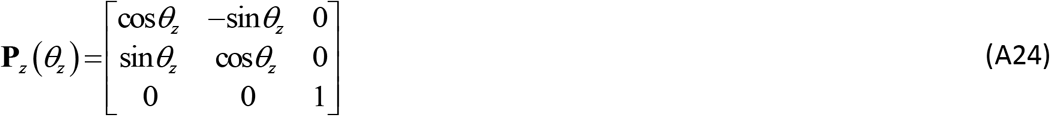

Here, we take the origin of the right- and left-eye coordinate systems to be the nodal point for that eye. Thus, a point **x** =(*x*, *y*,*z*) in the cyclopean space of Figure 2 would correspond to the point **x′** in the coordinate frame of the rotated right eye given by

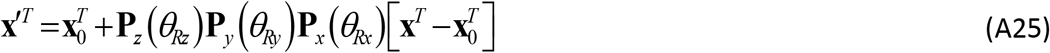

where **x**_0_ =(a,0,0).

The binocular matching equations for general eye rotations are given by first mapping the location in the right eye image plane into the cyclopean space (i.e., applying the operations in Equation A25 in the opposite order):

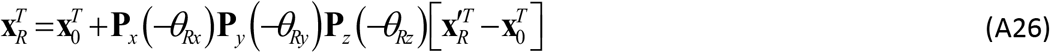

where 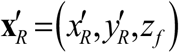. Next, the equations in the middle panel of Figure A1 are used to back project **x**_*R*_ =(*x*_*R*_, *y*_*R*_,*z*_*R*_) into scene space x (note that, in general, *z*_*R*_ ≠ *z*_*f*_). Then, the scene point is transformed into the left-eye coordinate frame:

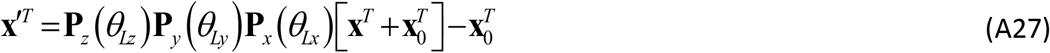

Finally, the scene point is projected to a point 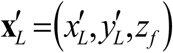 in the left-eye image plane, where 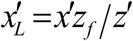 and 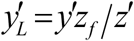 . This left eye point 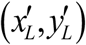 corresponds to the right eye point 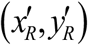, given the assumed slant, tilt and distance. Note that these matching left eye and right eye points can also be expressed in spherical coordinates (Equations A18–A21, with *a* set to zero).

Of course, the choice of coordinate system is arbitrary, in the sense that the matching process returns exactly the same information (the same slant, tilt and distance estimates) in all cases, given that the eye positions are known. However, the vision science literature uses many different coordinate systems to describe binocular geometry. The same terms can have different meanings depending on the choice of coordinate system. The possibility for confusion is high. For example, in image-plane coordinates when eyes are earth horizontal and in primary position (c.f. Figure 2) corresponding points always have the same y-coordinates; that is, there are no image-plane vertical disparities (see Figures 3B and 3C). On the other hand, in spherical coordinates, corresponding points have different elevation angles, and hence have angular vertical disparities. The important point here is that regardless of the image coordinate system chosen, the present matching process properly uses all of the stereo-geometry information for locally planar surfaces.

## References

Allenmark, F., & Read, J. C. (2011). Spatial stereoresolution for depth corrugations may be set in primary visual cortex. PLoS Computational Biology, 7(8), e1002142.

Backus, B. T., Banks, M. S., van Ee, R., & Crowell, J. A. (1999). Horizontal and vertical disparity, eye position, and stereoscopic slant perception. Vision Research, 39, 1143–1170.

Badcock, D.R., & Schor C.M. (1985). Depth-increment detection function for individual spatial channels. Journal of the Optical Society of America A, 2, 1211–1216.

Banks, M. S., Gepshtein, S., & Landy, M. S. (2004). Why is spatial stereoresolution so low? Journal of Neuroscience, 24, 2077–2089.

Blakemore, C. (1970a). A new kind of stereoscopic vision. Vision research, 10(11), 1181–1199.

Blakemore, C. (1970b). The range and scope of binocular depth discrimination in man. The Journal of Physiology, 211(3), 599–622.

Bohr I., & Read J.C. (2013). Stereoacuity with Frisby and revised FD2 stereo tests. PLoS One, 8(12), e82999.

Brainard, D. H. (1997). The Psychophysics Toolbox, Spatial Vision 10:433–436.

Bridge, H., & Cumming, B. G. (2001). Response of macaque V1 neurons to binocular orientation differences. Journal of Neuroscience, 21(18), 7293–7302.

Bridge, H., & Cumming, B. G. (2008). Representation of binocular surfaces by cortical neurons. Current Opinion in Neurobiology, 18(4), 425–430.

Bridge, H., Cumming, B. G., & Parker, A. J. (2001). Modeling V1 neuronal responses to orientation disparity. Visual Neuroscience, 18(6), 879–891.

Burge, J. (2020). Image-computable ideal observers for tasks with natural stimuli. Annual Review of Vision Science, 6: 491–517.

Burge, J., & Geisler, W.S. (2014). Optimal disparity estimation in natural stereo images. Journal of Vision, 14(2):1, 1–18.

Burge, J., Girshick, A.R., Banks, M.S. (2010). Visual-haptic adaptation is determined by relative reliability. Journal of Neuroscience, 30(22): 7714–7721.

Burge, J., McCann, B.C., Geisler, W.S. (2016). Estimating 3D tilt from local image cues in natural scenes. Journal of Vision, 16: 13, 1–25 doi:10.1167/16.13.2

Cagenello, R., & Rogers, B.J. (1993). Anisotropies in the perception of stereo surfaces: the role of orientation disparity. Vision Research, 33, 2189–2201.

Clark, J. J., & Yuille, A. L. (1990). Data Fusion for Sensory Information Processing Systems. Kluwer Academic Publishers.

Cochran, W.G. (1937). Problems arising in the analysis of a series of similar experiments. Supplement to the Journal of the Royal Statistical Society, 4, 102–118.

Cormack, L. K., Stevenson, S. B., & Schor, C. M. (1991) Interocular correlation, luminance contrast and cyclopean processing. Vision Research, 31, 2195–2207.

Coutant, B.E., & Westheimer, G. (1993). Population distribution of stereoscopic ability. Ophthalmology and Physiological Optics, 13, 3–7.

Cumming, B.G., & DeAngelis, G.C. (2001). The physiology of stereopsis. Annual Review of Neuroscience, 24, 203–238.

DeAngelis, G.C., Ohzawa, I., & Freeman, R.D. (1991). Depth is encoded in the visual cortex by a specialized receptive field structure. Nature, 352, 156–159.

Filippini, H. R., & Banks, M. S. (2009). Limits of stereopsis explained by local cross-correlation. Journal of Vision, 9(1):8, 1–18.

Fiorentini, A, & Maffei, L. (1971). Binocular depth perception without geometric cues. Vision Research, 11, 1299–1305.

Geisler W. S. (2011). Contributions of ideal observer theory to vision research. Vision Research, 51, 771–781.

Gillam, B., & Rogers, B. (1991). Orientation disparity, deformation, and stereoscopic slant perception. Perception, 20, 441–448.

Girshick, A.R., & Banks, M.S. (2009). Probabilistic combination of slant information: Weighted averaging and robustness as optimal percepts. Journal of Vision, 9(9):8, 1–20. doi:https://doi.org/10.1167/9.9.8

Goutcher, R., & Hibbard, P. B. (2014). Mechanisms for similarity matching in disparity measurement. Frontiers in Psychology, 4, 1014.

Green, D.M., & Swets, J.A. (1966). Signal Detection Theory and Psychophysics. New York: Wiley.

Greenwald, H. S., & Knill, D. C. (2009). Orientation disparity: a cue for 3D orientation? Neural Computation, 21(9), 2581–2604.

Halpern, D.L., Wilson, H.R., & Blake, R. (1996). Stereopsis from interocular spatial frequency differences is not robust. Vision Research, 36, 2263–2270.

Hibbard, P. B., & Langley, K. (1998). Plaid slant and inclination thresholds can be predicted from components. Vision Research, 38(8), 1073–1084.

Hibbard, P. B., Bradshaw, M. F., Langley, K., & Rogers, B. J. (2002). The stereoscopic anisotropy: Individual differences and underlying mechanisms. Journal of Experimental Psychology: Human Perception and Performance, 28(2), 469.

Hillis, J. M., Watt, S. J., Landy, M. S., & Banks, M. S. (2004). Slant from texture and disparity cues: optimal cue combination. Journal of Vision, 4(12), 967–992. http://doi.org/10.1167/4.12.1

Hirschmuller, H., & Scharstein, D. (2007). Evaluation of cost functions for stereo matching. IEEE Conference on Computer Vision and Pattern Recognition, 1–8.

Howard, I.P., & Rogers, B.J. (2012). Perceiving in Depth Vol. 2: Stereoscopic Vision. Oxford: Oxford University Press.

Jones, D.G., Malik, J. (1992). Determining three-dimensional shape from orientation and spatial frequency disparities. Proceedings of the European Conference on Computer Vision, Genova, 661–669.

Kato D, Baba M, Sasaki KS, & Ohzawa I. (2016). Effects of generalized pooling on binocular disparity selectivity of neurons in the early visual cortex. Philosophical Transactions of the Royal Society B, 371, 20150266.

Kim, S., Burge, J. (2018). The lawful imprecision of human surface tilt estimation in natural scenes. eLife, 7:e31148

Kim, S., Burge, J. (2020). Natural scene statistics predict how humans pool information across space in surface tilt estimation. PLoS Computational Biology, 16 (6), e1007947

Knill, D.C. (1998). Surface orientation from texture:ll ideal observers, generic observers, and the information content of texture cues. Vision Research, 38, 1655–1682.i

Knill, D. C., & Saunders, J. A. (2003). Do humans optimally integrate stereo and texture information for judgments of surface slant? Vision Research, 43(24), 2539–2558. http://doi.org/10.1016/S0042-6989(03)00458-9

Li, G., & Zucker, S.W. (2010). Differential geometric inference in surface stereo, IEEE Transactions on Pattern Analysis and Machine Intelligence, 32, 72–86.

Mitchison, G.J., & McKee, S.P. (1990). Mechanisms underlying the anisotopy of stereoscopic tilt perception. Vision Research, 30, 1781–1791.

Ogale, A.S., & Aloimonos, Y. (2005). Shape and the stereo correspondence problem. International Journal of Computer Vision 65(3), 147–162.

Ogle, K.N. (1938). Induced size effect: I. A new phenomenon in binocular space perception associated with the relative sizes of the images of the two eyes. Archives of Ophthalmology, 20(4), 604–623

Ohzawa, I., DeAngelis, G. C., & Freeman, R. D. (1990). Stereoscopic depth discrimination in the visual cortex: neurons ideally suited as disparity detectors. Science, 249(4972), 1037–1041.

Ohzawa, I., DeAngelis, G.C., & Freeman, R. D. (1997). Encoding of binocular disparity by complex cells in the cat’s visual cortex. Journal of Neurophysiology, 77, 2879–2909.

Oluk, C., & Geisler, W. S. (2020). Ideal Observers for the estimation of disparity in random-pixel stereograms. Journal of Vision, 20(11), 578–578.

Oruc, I., Maloney, L.T., & Landy, M.S. (2003). Weighted linear cue combination with possibly correlated error. Vision Research, 43, 2451–2468.

Pelli, D. G. (1997). The VideoToolbox software for visual psychophysics: Transforming numbers into movies, Spatial Vision 10:437–442.

Sanada, T. M., & Ohzawa, I. (2006). Encoding of three-dimensional surface slant in cat visual areas 17 and 18. Journal of neurophysiology, 95(5), 2768–2786.

Scharstein, D., & Szeliski, R. (2002). A taxonomy and evaluation of dense two-frame stereo correspondence algorithms. International Journal of Computer Vision, 47(1-3), 7–42.

Schumer, R.A., & Julesz, B. (1984). Binocular disparity modulation sensitivity to disparities offset from the plane of fixation. Vision Research, 24, 533–542.

Stevenson, S.B., Cormack, L.K., Schor, C.M. & Tyler, C.W. (1992). Disparity tuning in mechanisms of human stereopsis. Vision Research, 32, 1685–1694.

Stevens, K. A. (1983). Surface tilt (the direction of slant): a neglected psychophysical variable. Perception & Psychophysics, 33(3), 241–250. http://doi.org/10.3758/BF03202860

Super, B.J., & Klarquist, W.N. (1997). Patch-based stereo in a general binocular viewing geometry. IEEE Transactions on Pattern Analysis And Machine Intelligence, 19, 247–252.

Tyler, C.W., & Julesz, B. (1978). Binocular cross-correlation in time and space. Vision Research, 18, 101–105.

Vadal-Naguet, M. & Gepshtein, S. (2012). Spatially invariant computations in stereoscopic vision. Frontiers of Computational Neuroscience, https://doi.org/10.3389/fncom.2012.00047

Weiss Y., Simoncelli, E.P. & Adelson, E.H. (2002). Motion illusions as optimal percepts. Nature Neuroscience, 5, 598–604.

Wheatstone, C. (1838). On some remarkable, and hitherto unobserved, phenomena of binocular vision. Philosophical Transactions of the Royal Society of London, 128, 371–394

Wildes, R.P. (1991). Direct recovery of three-dimensional scene geometry from binocular stereo disparity. IEEE Transactions on Pattern Analysis and Machine Intelligence, 13, 761–774.

Wilson, H.R. (1976). The significance of frequency gradients in binocular grating perception. Vision Research, 16, 983–989.

